# Zinc excess promotes lysosome remodeling by activating HLH-30/TFEB through the action of the high zinc sensor HIZR-1

**DOI:** 10.64898/2026.02.22.707197

**Authors:** Ciro Cubillas, Hanwenheng Liu, Krupa Deshmukh, Adelita Mendoza, Daniel L. Schneider, Daniel Herrera, Chen Zhao, John T. Murphy, John Edwards, Abhinav Diwan, Kerry Kornfeld

## Abstract

Zinc is an essential element that plays many roles in animals. Since excess zinc is toxic, animals have evolved sophisticated mechanisms to achieve homeostasis. A conserved mechanism is to store excess zinc in specialized lysosomes, which contributes to zinc detoxification and provides a supply when zinc becomes limiting. In *C. elegans* excess zinc conditions promote an increase in lysosome-related organelles called bilobed granules. Defining how zinc-regulated transcription drives the observed lysosome remodeling is key to understanding zinc homeostasis processes. Here we describe a positively regulated zinc cascade controlled by *hizr-1* and *hlh-30/TFEB*, which encode the *C. elegans* high zinc sensor and the master regulator of the autophagy-lysosomal pathway, respectively; essential for connecting excess zinc homeostasis to lysosome biogenesis and remodeling. Our transcriptomic, genetic, and bioinformatic studies indicate that *hizr-1* and *hlh-30* are necessary and sufficient to activate transcription of lysosome genes under excess zinc conditions. In our regulatory model, zinc binding activates HIZR-1 protein, which accumulates in the nucleus and activates genes containing the HZA enhancer, including *hlh-30*. HLH-30 protein accumulates in the nucleus and activates genes containing the E-box enhancer. Genetic analysis of loss-of-function mutants demonstrated that both *hizr-1* and *hlh-30* are necessary for animals to tolerate excess zinc. Furthermore, HLH-30 promotes the increase in the number of acidified compartments, whereas HIZR-1 promotes the increase in the volume of the expansion compartment. These results define a genetic pathway that responds to excess zinc by increasing the number of lysosome-related organelles and their capacity to store and detoxify cytosolic zinc.

## Introduction

Zinc is an essential metal nutrient with a wide range of roles in biological systems. Zinc promotes catalysis when bound to enzyme active sites and the tertiary structure of proteins when bound to motifs such as zinc fingers in transcription factors; it has been experimentally estimated that around 10% of the human proteome binds zinc through cysteine interactions (Klug and Schwabe 1995; Auld and Bergman 2008; Andreini and Bertini 2012). In metazoans, zinc imbalances can lead to hormone-related and neurodegenerative diseases (Frederickson et al. 2004; Friedlich et al. 2004; Kodirov et al. 2006). Local and transient increasing zinc concentrations have been observed in multiple cellular contexts, including synapses (Tóth 2011), zinc-insulin co-release in pancreatic islets (Suckale and Solimena 2010), zinc sparks in early developmental stages of the embryo (Duncan et al. 2016; Zhang et al. 2016), during germline maturation (Mendoza et al. 2017; Zhao et al. 2018; Mendoza et al. 2022) and B cell development (Anzilotti et al. 2019); thus, these increases have a great impact on cell physiology and differentiation. In addition, intracellular zinc concentrations also rise in disease; it was found that ZIP14 overexpression in metastatic cancer increases zinc uptake leading to muscle loss by repressing MyoD and Mef2c expression (Wang et al. 2018) while studies revealed that ZIP8 upregulation in osteoarthritis cartilage of vertebrates increases intracellular zinc levels, leading to cartilage destruction by increasing metalloprotease gene expression (Kim et al. 2014). These observations indicate that changes in intracellular zinc significantly impact cell metabolism and development through the zinc mediated transcriptional response. Excess zinc is toxic (Maret and Sandstead 2006; Sandstead 2015; Schoofs et al. 2024); thus, organisms have evolved mechanisms to achieve homeostasis. Here we focus on how the transcriptional response that occurs during excess zinc is coordinated to achieve homeostasis.

The critical step in the response to zinc excess is zinc sensing. The zinc mediated signaling in animals is known to depend on transcription factors. In vertebrates and flies, the zinc-finger transcription factor MTF-1 mediates transcriptional activation of zinc transporters involved in zinc efflux that relieve zinc toxicity (Giedroc et al. 2001; Zhang et al. 2001; Yepiskoposyan et al. 2006; Hardyman et al. 2016). The ZNF658 zinc-finger transcriptional activator recognizes a DNA sequence called the zinc transcriptional regulatory element (ZRTE) located on the promoters of human genes coding for the ZnT5 and ZnT10 zinc transporters (Ogo et al. 2015). None of these transcription factors have been related to lysosome biogenesis or remodeling. In the nematode *C. elegans*, HIZR-1 is the high zinc sensor and master regulator of high zinc homeostasis (Dietrich et al. 2016; Warnhoff et al. 2017; Earley, Mendoza, et al. 2021). HIZR-1 is a nuclear receptor transcription factor that directly binds zinc in the ligand-binding domain. Zinc binding triggers HIZR-1 to accumulate in the nucleus where it directly binds the high zinc activation (HZA) element, an enhancer that is present in the promoters of multiple genes that are activated by excess zinc (Roh et al. 2015; Liu et al. 2025). The known transcriptional response to excess zinc in *C. elegans* primarily occurs in the intestine, and the promoters of zinc-activated target genes contain a GATA motif driving intestinal expression in addition to the HZA enhancer (Roh et al. 2015). Target genes that are activated by excess zinc promote detoxification by three basic mechanisms. (1) Activation of metallothionein-coding genes *mtl-1* and *mtl-2* promotes zinc binding and sequestration in the cytosol (Roh et al. 2015; Warnhoff et al. 2017). (2) Activation of the zinc transporter gene *ttm-1* promotes excretion of zinc from the apical surface of intestinal cells (Roh et al. 2013). (3) Activation of the zinc transporter gene *cdf-2* promotes zinc transport into the lumen of lysosome-related organelles (LRO), which both store and detoxify excess zinc (Davis et al. 2009; Roh et al. 2012). In response to excess zinc, there is an increase in the number of LROs, which increases the capacity for zinc storage (Shomer et al. 2019). Furthermore, LROs are remodeled in excess zinc (Roh et al. 2012). Mendoza et al (2024) showed that the LROs consist of an acidified compartment and an expansion compartment; the volume of the expansion compartment increases around six-fold in response to excess zinc (Mendoza et al. 2024). Zinc storage in lysosomes has been conserved during evolution. Mammals contain zinc-rich vesicles called zincosomes that are labelled by the dye FluoZin (Palmiter et al. 1996; Haase et al. 2008). A number of genetic pathways required for biogenesis of zinc-rich LROs have been described in *Drosophila melanogaster* (Tejeda-Guzmán et al. 2018) and *C. elegans* (Roh et al. 2012). In *C. elegans,* the *glo-3* and *pgp-2* genes are necessary for the assembly of functional bilobed granules, and loss-of-function mutants are hypersensitive to excess zinc (Roh et al. 2012); by contrast, CDF-2 is not necessary for LRO remodeling, but it is necessary to transport zinc into these LROs (Roh et al. 2012; Warnhoff et al. 2017). Thus, CDF-2 is an LRO resident protein that is transcriptionally regulated by *hizr-1*.

It is unknown how excess zinc sensed by *hizr-1* results in the increase in the number of LROs and the morphological change of the expansion compartment. To address these questions, we used genome-wide, genetic, and bioinformatic approaches to characterize the transcriptional response to excess zinc. Our results identify many new genes that are activated or repressed by excess zinc, and a significant number of activated genes require *hizr-1* for regulatory control. We focused on a group of genes that are part of the autophagy-lysosome pathway, and its regulator *hlh-30*/TFEB. In metazoans, lysosomal biogenesis is controlled by TFEB, which transcriptionally activates genes belonging to the Coordinated Lysosomal Expression and Regulation (CLEAR) network (Settembre et al. 2011). Expression of TFEB is self-regulated, and it positively regulates a set of target genes required for lysosome biogenesis through binding of a DNA element called the E-box. This promotes transcriptional activation of autophagy, longevity, stress response, starvation and innate immune responses, a regulatory control conserved in *C. elegans* (Lapierre et al. 2013; O’Rourke and Ruvkun 2013; Visvikis et al. 2014; Lin et al. 2018). Both excess zinc (Wei et al. 2018) and zinc deficient conditions (North et al. 2012; Kawamata et al. 2017) have been reported to transcriptionally activate autophagy in eukaryotes; although there is evidence supporting the idea that zinc acts as a positive regulator of autophagy, the underlying mechanism has not been elucidated (Liuzzi et al. 2014).

Here we show that the *hlh-30* gene contains an HZA enhancer in its promoter and is a direct target gene of *hizr-1*. *hlh-30* transcripts accumulate in response to excess zinc, and HLH-30 protein translocates to the nucleus to regulate gene expression. This response is conserved in human cells, since our results indicate that TFEB accumulates in the nucleus in response to excess zinc. In *C. elegans*, *hlh-30* is necessary for tolerance to excess zinc, indicating it plays a functional role in homeostasis. By analyzing the genome-wide response to excess zinc, we identified target genes having both an HZA enhancer and an E-box enhancer and requiring both HIZR-1 and HLH-30 for full activation. These findings define a gene regulatory cascade with *hizr-1* activating *hlh-30,* and these two transcription factors functioning together and separately to control gene expression. We used super-resolution microscopy to determine the role of these genes in LRO remodeling. Our results indicate that *hlh-30* is necessary to increase the number of acidified compartments in response to excess zinc, whereas *hizr-1* is necessary to generate CDF-2 positive vesicles that increase the volume of the expansion compartment. Together, these results identify a pathway that plays a central role in high zinc homeostasis by increasing the number of LROs that can accommodate excess zinc and driving their morphological remodeling.

## Results

### Identification of genes regulated by excess dietary zinc in wild-type *C. elegans*

The main experimental variables affecting/influencing transcriptional response to excess dietary zinc in *C. elegans* are the concentration of zinc, duration of exposure, and the developmental stage of animals. To characterize the genome-wide transcriptional response to zinc excess, we cultured mixed-stage populations of wild-type *C. elegans* for 16-18h on Noble-Agar Minimal Medium (NAMM) dishes supplemented with 0μM zinc (zinc replete) or 200μM zinc (zinc excess). Cultures included a lawn of *E. coli*, which contributes an undefined amount of zinc to the diet. RNA was purified and analyzed by RNA sequencing (RNA-seq), establishing an expression value for each detectable gene in the two conditions (20,148 genes, Table S1). Known zinc-activated genes, such as *hizr-1*, *cdf-2, ttm-1, mtl-1,* and *mtl-2,* and known zinc repressed genes, such as *zipt-2.3,* displayed the expected regulation, indicating the experiment was successful (Figure 1A) (Davis et al. 2009; Roh et al. 2013; Roh et al. 2015; Warnhoff et al. 2017; Earley, Cubillas, et al. 2021; Mendoza et al. 2024). In zinc excess, 967 and 457 genes displayed significantly higher and lower levels of transcripts (*P* ≤ 0.001), respectively. The magnitude of activation ranged from 0.45 to 6.95 log2 fold change, whereas the magnitude of repression varied from -0.32 to -5.01 log2 fold change (Table S2).

**Figure 1.**
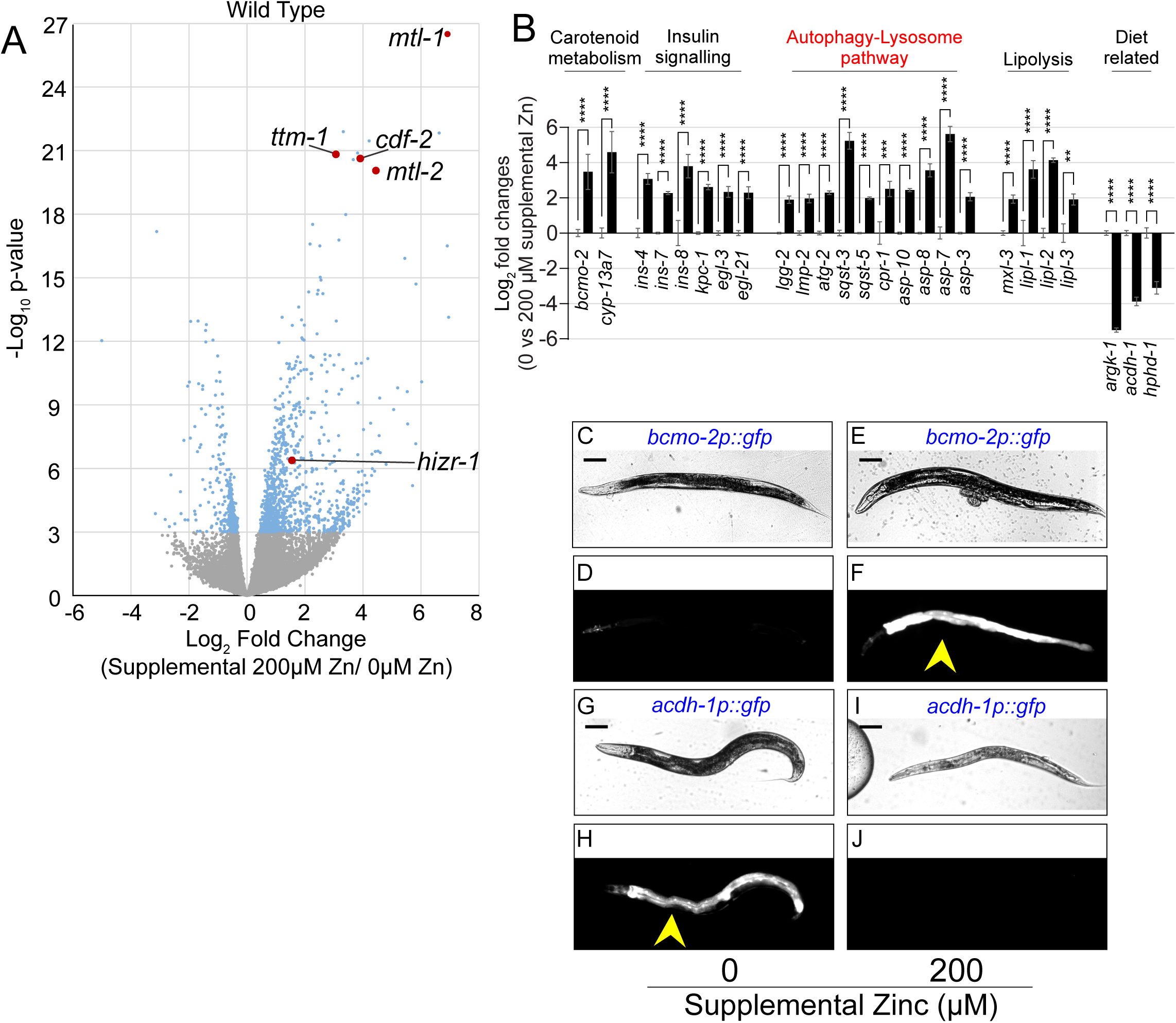
Identification and characterization of zinc-regulated genes in wild type. (A,B) Mixed-stage populations of WT worms were cultured on zinc replete (0μM supplemental ZnSO_4_) or zinc excess (200μM supplemental ZnSO_4_) NAMM dishes for 16-18 hours, and gene expression was analyzed. (A) RNA sequencing. Each point is data for one gene, showing Log_2_ fold change of expression level and the *P-*value. Blue and gray indicate significant and nonsignificant *P*-values, respectively. Red indicates genes previously established to be zinc-activated. (B) Gene expression was analyzed by qRT-PCR. Bars indicate the Log_2_ fold change of mRNA levels – the level in replete conditions (0μM supplemental zinc) was set equal to 0 and compared to the value at 200μM supplemental zinc. Statistical analysis is done on at least three independent experiments, using one way ANOVA with post-hoc Dunnett T3. For significant p values: *<0.05; **<0.01; ***<0.001; ****<0.0001. Non-significant p values are indicated as “ns”. Error bars represent mean ± SD. Gene names are below, and pathway names are above. (C-J) Transgenic animals containing plasmids with the *bcmo-2* promoter or *acdh-1* promoter driving GFP were cultured on NAMM dishes with replete zinc (0μM supplemental ZnSO_4_) or zinc excess (200μM supplemental ZnSO_4_) for 16-24 h and visualized by bright field (upper) or epifluorescence (lower) microscopy. Yellow arrows indicate fluorescence in the intestine. Scale bar = 100µm. *bcmo-2* was activated by zinc excess, whereas *acdh-1* was repressed by zinc excess. Images are representative of at least 2 independent experiments.

As a complementary approach, we performed a conceptually similar experiment with different medium conditions, duration of exposure, stage of animals, and detection method for RNA abundance. We used fully defined liquid *C. elegans* maintenance medium (CeMM) containing 30μM zinc (zinc replete) or 500μM zinc (zinc excess) (Davis et al. 2009). Animals were cultured for at least 6 days in these conditions to achieve steady state gene expression, RNA was isolated from synchronized L4 stage animals, and transcript levels were determined by using DNA microarrays, establishing an expression value for each detectable gene in the two conditions based on four biological replicates (12,396 genes, Table S3). Known zinc-activated genes, such as *cdf-2, ttm-1, mtl-1,* and *mtl-2,* displayed the expected regulation, indicating the experiment was successful (Figure S1). In zinc excess conditions, 134 and 113 genes displayed significantly higher and lower levels of transcripts, respectively (P ≤ 0.01). The magnitude of activation ranged from 0.06 to 3.95 log_2_ fold change, whereas the magnitude of repression varied from -0.06 to -4.01 log_2_ fold change (Table S4). Comparing the results of microarray and RNAseq experiments revealed that of the 134 activated genes, 46 (35%) were identified by both methods (Figure S2); of the 113 repressed genes, 4 (4%) were shared by both methods (Figure S2). We found *mtl-1, mtl-2, cdf-2,* and *ttm-1* were activated by excess zinc as measured by RNAseq and microarrays, suggesting that some genes are robustly regulated in a variety of conditions, whereas other genes may be regulated only under specific conditions.

To validate transcript levels determined by RNA-seq or microarray, we performed quantitative real-time reverse transcription polymerase chain reaction (qRT-PCR) using independently derived RNA from mixed-stage populations of wild-type *C. elegans* cultured with 0μM zinc (zinc replete) or 200μM zinc (zinc excess). Tables S5 and S6 show158 zinc regulated genes, including 133 zinc-activated genes and 25 zinc-repressed genes that were validated by this independent method.

To analyze the specificity of induction by zinc, we determined how eleven zinc-regulated genes responded to excess manganese, a physiological metal that is also a divalent cation. We selected seven genes that range from strongly to moderately activated and four genes that range from strongly to moderately repressed. Mixed-stage populations of wild-type *C. elegans* were cultured for 16-18h on Noble-Agar Minimal Medium (NAMM) dishes supplemented with 200μM zinc (zinc excess) or 500μM manganese (manganese excess). RNA was purified and analyzed by qRT-PCR. Seven genes were significantly more activated in excess zinc compared to excess manganese excess. The four genes that were significantly repressed by excess zinc were not significantly repressed by excess manganese (Figure S3, Table S7). Thus, the response to excess zinc does not appear to be a nonspecific response to any excess divalent cation.

We identified multiple metabolic pathways that are regulated by excess zinc. The genes *bcmo-2, cyp-13a4, cyp-13a5 and cyp-13a7* encode known components of the retinoic acid/carotenoid metabolism pathway and have also been found to be activated by excess cadmium (Cui and Freedman 2009; Earley, Cubillas, et al. 2021). The insulin and neuropeptide signaling and processing genes were represented by *ins-4, ins-7,* and *ins-8* (Ritter et al. 2013; Zhu and Chin-Sang 2024) encoding for insulin and other neuropeptides as well as their predicted convertase coding genes *kpc-1, elg-3 and egl-21* (Stawicki et al. 2013; Li and Kim 2014; Nkambeu et al. 2019; Zhu and Chin-Sang 2024). The autophagy-lysosome pathway had the highest number of zinc activated genes; we identified at least 22 genes, including multiple that encode aspartases (*asp-* genes) and proteins involved in autophagy such as *lgg-2, atg-2, sqst-1, sqst-3,* and *sqst-5;* from these, *sqst-5* has been reported to be involved in zinc homeostasis (Evans et al. 2020). We identified multiple genes involved in lipolysis, primarily the lipase genes *lipl-1, lipl-2,* and *lipl-3*, alongside *mxl-3*, coding for a transcription factor involved in the response to starvation (O’Rourke and Ruvkun 2013). Interestingly, a known heme responsive gene *hrg-7 (asp-10)* was found to be activated by zinc (Sinclair and Hamza 2010; Sinclair et al. 2017); meanwhile, genes involved in B12 metabolism such as *acdh-1* and *hphd-1* were found to be zinc repressed (Watson et al. 2016; Fox et al. 2022; Ponomarova et al. 2023). Genes in these pathways were confirmed to be activated by excess zinc by qRT-PCR, consistent with the RNAseq data (Figure 1B, Table S8).

We selected five zinc-activated genes (*bcmo-2, thn-2, cyp-14a4, ins-8, ugt-1*) and one zinc-repressed gene (*acdh-1*) to determine the localization of their gene expression *in vivo*. Plasmids containing predicted promoter sequences driving GFP expression were introduced into transgenic animals. L4 stage animals were cultured on NAMM dishes containing either 0 or 200μM supplemental zinc and imaged after 16-18h. All six genes displayed the predicted regulation; five were expressed primarily in the intestine, whereas *ins-8* was primarily expressed outside the intestine (Figure 1C-J, S4). These results are consistent with previous findings that zinc activated genes are primarily expressed in intestinal cells, including *hizr-1*, *cdf-2, mtl-1,* and *mtl-2* (Roh et al. 2015).

### The *hizr-1* transcription factor is necessary for many genes to be regulated by excess zinc

HIZR-1 is a nuclear receptor transcription factor that was identified in a forward genetic screen because it regulates activation of the *cdf-2* promoter by excess zinc (Warnhoff et al. 2017). *hizr-1* was demonstrated to be necessary for transcriptional activation of several other genes in excess zinc, including *ttm-1, mtl-1,* and *mtl-2*. Two classes of mutations were identified in the screen: 1) loss-of-function mutations (i.e. *hizr-1* Q87Stop) abrogate activation of *cdf-2:gfp* transcription in excess zinc; 2) a gain-of-function mutation (*hizr-1* D270N) causes high level expression of *cdf-2:gfp* transcripts in zinc replete medium. However, the genome-wide effects of *hizr-1* on zinc-regulated transcription were not defined; thus, we used RNA-seq to determine transcript levels in *hizr-1*(*am286 Q87Stop*) mutants. NAMM dishes were supplemented with either 0 or 200μM zinc, mixed stage animals were cultured for 16-18h, RNA was purified, and gene expression was analyzed by RNA-seq. We first compared *hizr-1(am286lf)* mutant gene-expression values in basal and excess zinc conditions (Figure 2A, Table S9). Many genes displayed lower levels of activation compared to wild type, including all five well-established zinc-activated genes. For some genes, activation was reduced but still detectable (*ttm-1*), whereas in some genes activation was abrogated (*mtl-2*) (Figure 2A).

**Figure 2.**
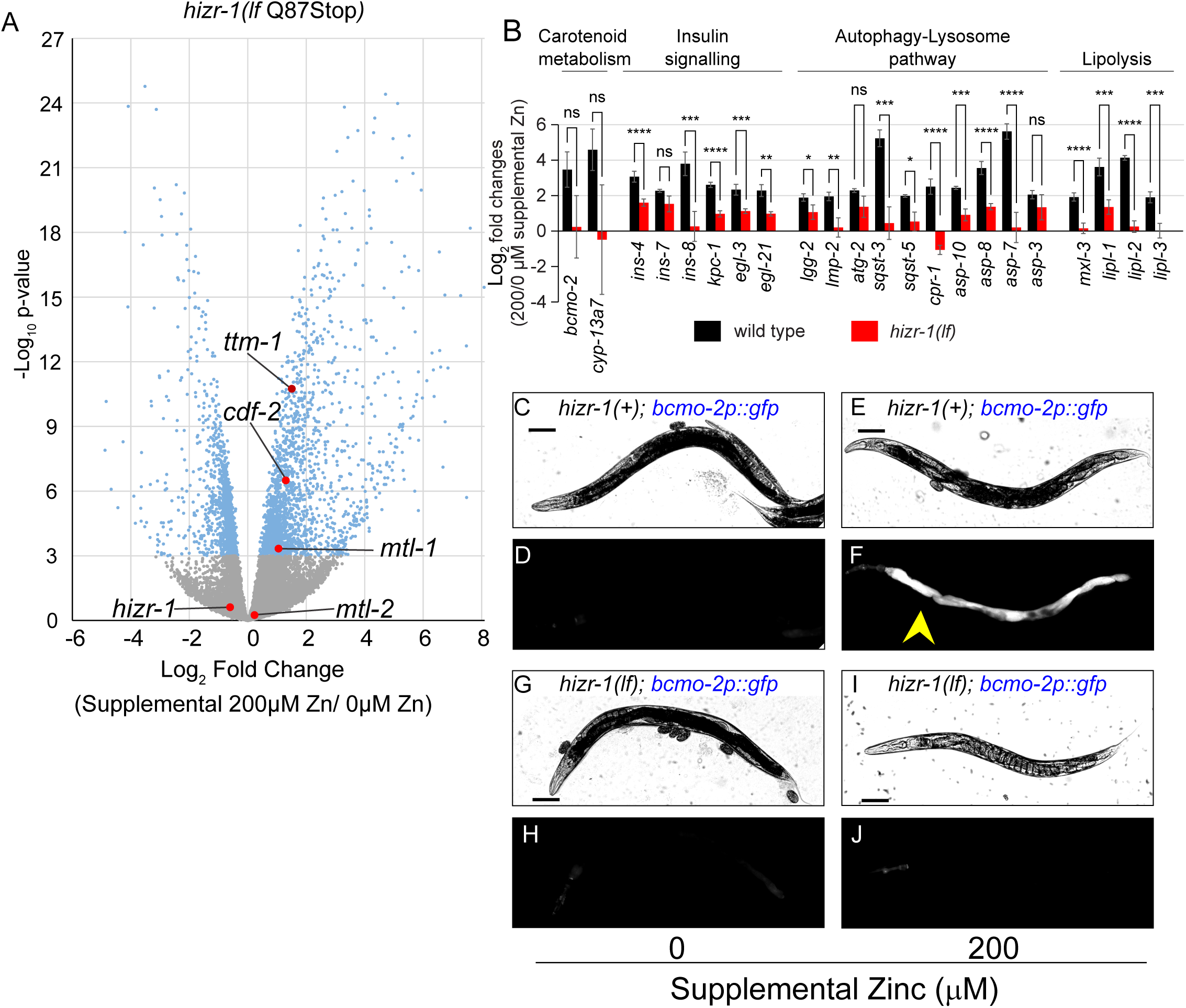
*hizr-1* is necessary for many genes to be activated by excess zinc. (A) Mixed-stage populations of *hizr-1(am286lf)* worms were cultured on zinc replete or zinc excess NAMM dishes for 16-24 h, and gene expression was analyzed by RNA sequencing. Each point is data for one gene, showing Log_2_ fold change of expression level and the *P*-value. Blue and gray indicate significant and nonsignificant *P*-values, respectively. Red indicates genes previously established to be zinc-activated in wild type. (B) Mixed-stage populations of WT and *hizr-1(am286lf)* worms were cultured on zinc replete or zinc excess NAMM dishes for 18-24 h, and gene expression was analyzed by qRT-PCR. Bars indicate the ratio of normalized mRNA levels in zinc excess/zinc replete conditions. Statistical approach is described in Figure 1 legend. Gene names are below, and pathway names are above. WT data are identical to Figure 1B but displayed in a different manner. (C-J) Transgenic animals containing plasmids with the *bcmo-2* promoter driving GFP were cultured on NAMM dishes with 0 or 200µM supplemental zinc for 16-24h and visualized by bright field (upper) or fluorescence (lower) microscopy. In panels G-J, the strain contains the *hizr-1(am286lf)* mutation. Scale bar = 100µm. *bcmo-2* was activated by zinc excess in the WT background (yellow arrow), but not in the *hizr-1(lf)* background.

To validate RNA-seq results with an independent method, we used qRT-PCR to analyze genes belonging to the four pathways activated by zinc excess. Wild type and *hizr-1(am286lf)* mutants were compared. Seventeen of those genes displayed significantly reduced or absent activation in *hizr-1* mutants (Figure 2B, Table S10); five genes displayed a trend towards reduced activation that was not significant with this sample size. Thus, *hizr-1* contributes to the transcriptional activation of many genes that are regulated by excess zinc. To extend these findings to localization *in vivo,* we used genetic crosses to combine the *bcmo-2p:gfp* marker with the *hizr-1(am286lf)* mutation. In excess zinc, *bcmo-2* is dramatically activated in intestinal cells of wild-type animals, but this activation was abrogated in the *hizr-1(am286lf)* mutant (Figure 2C-J).

### *hlh-30*, a regulator of the autophagy-lysosomal pathway, is activated by excess zinc and necessary for excess zinc homeostasis

The *hlh-30* gene encodes a helix-loop-helix transcription factor that binds to E-box enhancers in promoters and thereby controls multiple genes involved in autophagy and lysosome biogenesis (Ephrussi et al. 1985; Massari and Murre 2000; Lapierre et al. 2013). The *hlh-30* promoter contains an E-box, indicating that *hlh-30* positively regulates its own promoter (Grove et al. 2009; Lapierre et al. 2013). *hlh-30* has been implicated in a wide range of biological processes, including starvation response (Settembre et al. 2013; Murphy et al. 2019), aging (Lapierre et al. 2013; Lin et al. 2018), and innate immunity (Visvikis et al. 2014; Chen et al. 2017; Kumar et al. 2019), but it has not been previously implicated in zinc homeostasis or lysosome remodeling. Our discovery that multiple genes in the autophagy-lysosomal pathway are activated by excess zinc led us to hypothesize that *hlh-30* plays a role in this process. To investigate this hypothesis, we analyzed the *hlh-30* promoter for an HZA enhancer. Indeed, at position -235 relative to the translation start site, the *hlh-30* promoter contains a sequence that closely resembles established HZA enhancers (Figure 3A,B). If this sequence is a functional HZA enhancer, then we predict that *hlh-30* will be activated by excess zinc and *hizr-1* will be necessary for this activation. To directly evaluate regulation, we measured *hlh-30* mRNA levels by qRT-PCR in wild type and *hizr-1(am286lf)* mutant animals exposed to replete or excess zinc. In wild-type animals, *hlh-30* mRNA levels increased ∼5-fold in 200μM supplemental zinc compared to zinc replete conditions; *hlh-30* mRNA accumulation was significantly lower, ∼1.7-fold, in *hizr-1(am286lf)* mutants (Figure 3C, Table S11). Thus, *hlh-30* is activated by excess zinc, and *hizr-1* is necessary for full activation, suggesting HIZR-1 binds the HZA enhancer in the *hlh-30* promoter. To explore regulation of HLH-30 protein levels *in vivo*, we analyzed transgenic animals with multicopy extrachromosomal arrays expressing HLH-30 fused to a red fluorescent protein (*hlh-30p:*HLH-30*::rfp*) (Murphy et al. 2019). Synchronized animals were cultured with 0 or 200μM supplemental zinc for 16h and imaged by fluorescence microscopy (Figure 3D-G). We focused on the intestinal cells, since this is the site of excess zinc homeostasis and the known site for nuclear HLH-30 localization under standard growth conditions. The proportion of animals that displayed HLH-30 nuclear accumulation in intestinal cells was quantified. HLH-30 nuclear accumulation increased significantly from ∼3% in zinc replete conditions to ∼40% in excess zinc (Figure 3H). Thus, excess zinc causes accumulation of *hlh-30* mRNA and nuclear localization of HLH-30 protein, indicating HLH-30 is strongly activated by excess zinc.

**Figure 3.**
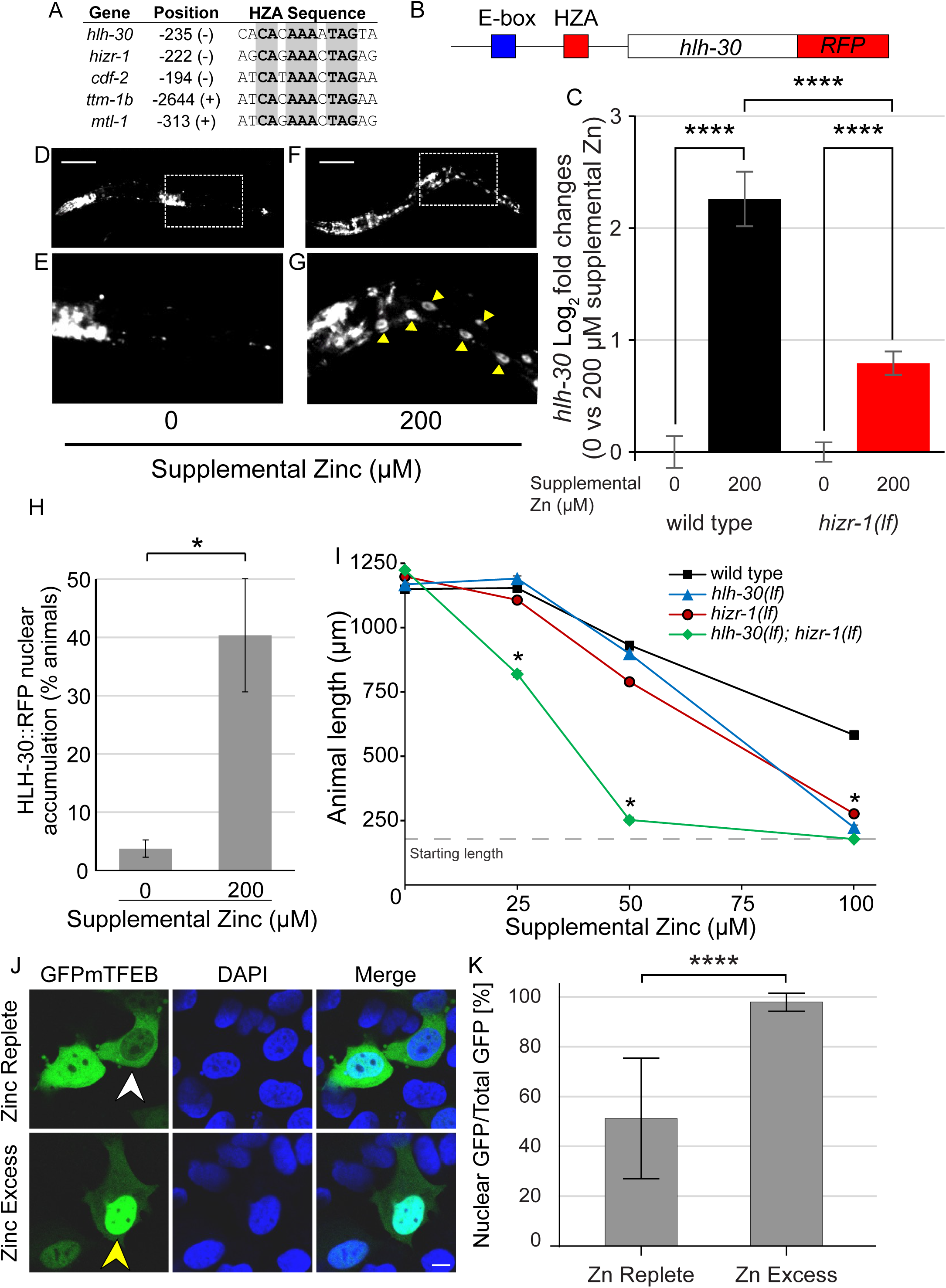
HLH-30 is activated by high zinc and necessary for high zinc homeostasis. (A) The alignment shows DNA sequences from the promoters of *hlh-30*, *hizr-1*, *cdf-2*, *ttm-1b*, and *mtl-1*. Bold and gray indicate nucleotides identical in all five sequences. Position (+) or (-) indicates the number of base pairs upstream of the translation start site and the orientation of the HZA sequence (Roh et al. 2015). (B) A diagram of a portion of the extrachromosomal array in the *hlh-30p::hlh-30::rfp* strain displaying the E-box (blue) and HZA (red) enhancers in the promoter (black line), the *hlh-30* coding region (open box), and the RFP coding region (red box). (C) Mixed-staged WT or *hizr-1(am286lf)* animals were cultured with 0 or 200μM supplemental zinc for 18-24 h, harvested for RNA, and analyzed by qRT-PCR. Bars indicate the Log_2_ fold change of mRNA levels – the level in replete conditions (0μM supplemental zinc) was set equal to 0 and compared to the value at 200μM supplemental zinc. Statistical approach is described in Figure 1 legend. Genotype and supplemental zinc are shown below. (D-G) Transgenic animals containing *hlh-30p*::*hlh-30*::*rfp* at the L2-3 stage were cultured with 200μM zinc for 16h. D,E show a fluorescence microscope image of one animal cultured with 0μM supplemental zinc – the dotted line box is enlarged in E. Nuclear localization of HLH-30 was not observed. Images are representative of at least 2 independent experiments. F,G show one animal cultured with 200μM supplemental zinc – the dotted line box is enlarged in G. Nuclear localization of HLH-30 is indicated by yellow arrowheads. Scale bars = 100µm. (H) Fluorescent images of individual animals were scored as positive or negative for nuclear accumulation of HLH-30::RFP in intestinal cells. Statistical approach is described in Figure 1 legend, but done with Welch’s t test. (I) Wild type, *hizr-1(am286lf)*, *hlh-30(tm1978lf)*, and *hlh-30(lf);hizr-1(lf)* double mutant hermaphrodites were synchronized at the L1 stage and cultured on NAMM dishes for 3 days with the indicated levels of supplemental zinc. The length of individual animals was measured using microscopy and ImageJ software. Statistical analysis by two-way anova. (J) Representative confocal images of HEK cells transfected with GFP-tagged mTFEB and treated for 4 hours with either normal medium (Zinc Replete) or medium supplemented with 200 μM ZnSO (Zinc Excess). GFP-mTFEB (green), DAPI (blue), and merged signals are shown. Arrowheads indicate representative cells exhibiting cytoplasmic (white, top) or nuclear (yellow, bottom) localization of TFEB. Scale bar =10 μm. (K) Quantification of nuclear GFP intensity as a percentage of total cellular GFP intensity under Zinc Replete (n = 86) and Zinc Excess (n = 76) conditions. Data are presented as mean ± SD. ****p < 0.0001 by Mann-Whitney test.

To determine the function of *hlh-30* in zinc homeostasis, we analyzed the *hlh-30(tm1978)* deletion mutation which is a strong loss-of-function or null allele (Visvikis et al. 2014; Chen et al. 2017). Length of animals after 3 days of growth in excess zinc was evaluated, since this is a sensitive and quantitative measure of homeostatic function. Wild-type animals displayed normal growth in 25μM supplemental zinc and a dose dependent reduction in growth rate in 50 and 100μM supplemental zinc (Figure 3I). *hlh-30(tm1978)* mutants displayed hypersensitivity to excess zinc, with significantly less growth in 100μM supplemental zinc compared to wild type (Figure 3I). This pattern is similar to *hizr-1(am286lf)* mutants, which were previously demonstrated to be hypersensitive to excess zinc (Figure 3I) (Warnhoff et al. 2017). Thus, *hlh-30* is necessary for robust growth in excess zinc. To evaluate the interaction between *hlh-30* and *hizr-1*, we constructed and analyzed a *hlh-30; hizr-1* double mutant. Double mutant animals displayed synergistic hypersensitivity to excess zinc; at 25 and 50μM supplemental zinc, *hizr-1* and *hlh-30* single mutants displayed growth similar to wild type, whereas the *hlh-30; hizr-1* double mutant displayed significantly reduced growth compared to wild type and the single mutants (Figure 3I). Thus, *hlh-30* and *hizr-1* play important and non-redundant roles in excess zinc homeostasis, since both single mutants displayed defects at high supplemental zinc. Since both genes encode transcription factors, non-redundant roles indicates these transcription factors may control some separate target genes. In addition, *hlh-30* and *hizr-1* appear to play redundant roles, revealed by the synergistic defects displayed by double mutants at moderate supplemental zinc. This finding suggests these two transcription factors may share some common target genes.

The HLH-30/TFEB roles have been highly conserved during evolution across species. To determine if mammalian TFEB is also activated by excess zinc, we analyzed mammalian cultured cells that express TFEB fused to GFP. In standard culture medium (zinc replete), about half the GFP fluorescence is localized to the cytoplasm and about half is localized in the nucleus. In response to excess zinc in the medium, there is a significant shift of HLH-30 into the nucleus, consistent with the model that activation by excess zinc is evolutionarily conserved (Figure 3J-K).

### A subset of HIZR-1 target genes contain an E-box, indicating these promoters are regulated by binding of HIZR-1 and HLH-30

The HIZR-1 nuclear receptor transcription factor directly binds the HZA enhancer (Warnhoff et al. 2017). In comparison, the HLH-30 helix-loop-helix transcription factor directly binds the E-box enhancer (Massari and Murre 2000; Grove et al. 2009; Palmieri et al. 2011). The finding that the *hlh-30* promoter contains both an HZA and an E-box enhancer suggests this promoter is coordinately controlled by both transcription factors and suggests promoters from other zinc activated genes may also contain both enhancers. To test this prediction, we analyzed the predicted promoters of 106 zinc-activated genes that require *hizr-1* for full activation to determine if they contain an HZA, an E box, or both enhancers. To define promoter regions, we used RSAT (Medina-Rivera et al. 2015) to collect the upstream sequence from each candidate gene, beginning from the predicted start ATG codon and extending to 2000bp upstream. To search for enhancers, we defined the HZA (Roh et al. 2015) element and the E-box (Weirauch et al. 2014) based on reported weight matrices. We searched the promoter regions for matches to the HZA and the E-box weight matrices simultaneously (Figure 4A) by using the *cis*-Regulatory element enriched Regions (CRERs) program also available at RSAT (Turatsinze et al. 2008; Medina-Rivera et al. 2015). This analysis revealed that all 106 promoters contained a sequence similar to the HZA and defined two classes of promoters: class H promoters have a predicted HZA but not a predicted E-box; class HE promoters have a predicted HZA and a predicted E-box within a 50bp interval or longer (Figure 4A-C, Table S12). One of the caveats of this analysis is that it may have excluded relevant promoter sequences that are further upstream of the cut off or downstream of the ATG; however, we think the experimental definition of the promoter region is reasonable given the typically small size of *C. elegans* genes. A weight matrix is a probabilistic description of an enhancer sequence, and the analysis may have scored as negatives those sequences that are weak matches but nonetheless functional *in vivo* or scored as positives those sequences that may not function as an enhancer *in vivo*. For example, the *hizr-1* promoter contains a sequence similar to the E-box weight matrix at position -563; however, a *hizr-1* promoter driving GFP was strongly activated by excess zinc in *hlh-30(lf)* animals, indicating *hlh-30* is not necessary to regulate *hizr-1* in excess zinc (Figure S5).

**Figure 4.**
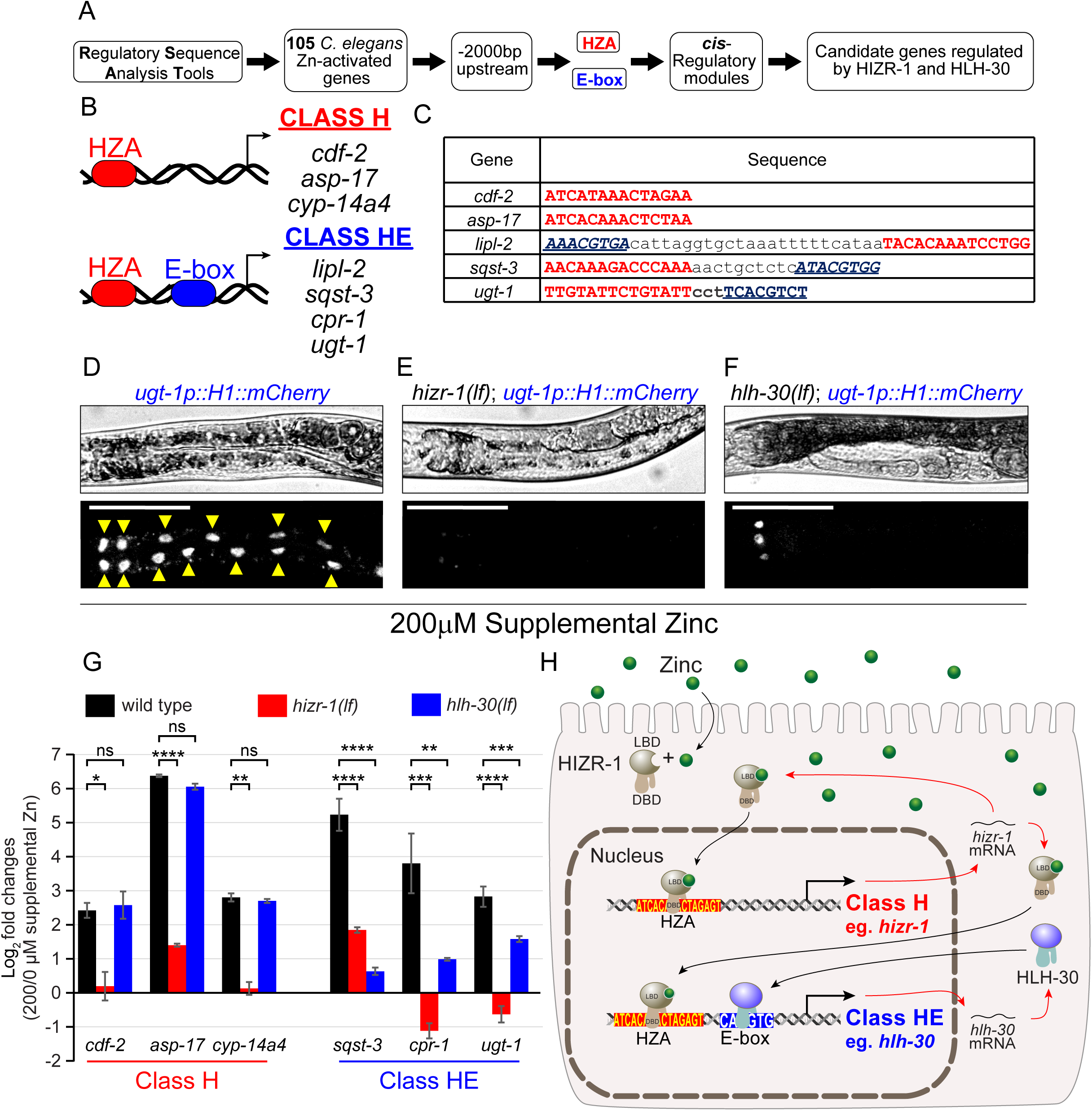
HIZR-1 and HLH-30 coordinately regulate class HE target genes. (A) An overview of the strategy to identify candidate genes regulated by HIZR-1 and HLH-30. The Regulatory Sequence Analysis Tools database was used to obtain the predicted promoters of zinc-activated genes identified in this study. MEME was used to generate an updated HZA weight matrix based on 17 *C. elegans* zinc-activated genes instead of only 4 genes as previously described (Roh et al. 2015). We used the previously defined E-box weight matrix for HLH-30 (Grove et al. 2009). The Cis-Regulatory element Enriched Regions program was used to search the promoter regions for matches to the HZA and E-box weight matrices within a 50bp distance, generating a list of genes that are candidates for requiring HIZR-1 and HLH-30 for full activation in excess zinc (Table S12). (B,C) Cartoons show DNA and transcription start site (arrow) for class H (*cdf-2*, *asp-17, cyp-14a4*) and class HE (*lipl-2*, *sqst-3, cpr-1, ugt-1*) genes. Table shows sequence of HZA (red), E-box (blue), and intervening sequences (black). (D-F) Transgenic animals containing plasmids with the *ugt-1* promoter driving H1::mCherry were cultured on NAMM dishes with 200μM supplemental zinc for 16-24h and visualized by bright field (upper) or fluorescence (lower) microscopy. Yellow arrows indicate fluorescence in intestinal nuclei of the nuclear localized H1::mCherry. Scale bar = 100 μm. *ugt-1* was activated by excess zinc in wild type, and *hizr-1* and *hlh-30* were both necessary for full activation. (G) Mixed staged wild type, *hizr-1(am286lf)*, and *hlh-30(tm1978lf)* populations of synchronized animals were cultured with 0 or 200μM supplemental zinc for 16-18h. Gene expression was analyzed by qRT-PCR. Gene and class names are below. Bars indicate the ratio of normalized mRNA levels in zinc excess/zinc replete conditions. Statistical approach is described in Figure 1 legend. (H) A cartoon showing HIZR-1 binding zinc (green circle) in the ligand-binding domain (LBD), entering the nucleus (black arrow), and binding HZA enhancers via the DNA-binding domain (DBD). HIZR-1 binds its own promoter, resulting in auto-activation (red arrow), the promoter of *hlh-30*, a class HE gene, as well as other class H and class HE genes. HLH-30 binds its own promoter, resulting in auto-activation (blue arrow), as well as other class HE genes.

We predicted that class H genes would require *hizr-1* but not *hlh-30* for zinc-activated transcription, whereas class HE genes would require both *hizr-1* and *hlh-30* for full activation. To test this prediction, we used the method of qRT-PCR to analyze wild type, *hizr-1(am286lf)*, and *hlh-30(tm1978lf)* mutant animals. The activation of the class H genes *cdf-2, asp-17*, and *cyp-14a4* by excess zinc was significantly reduced in *hizr-1(lf)* mutants but was not significantly affected in *hlh-30(lf)* mutants (Figure 4G, Table S13). By contrast, activation of the class HE genes *sqst-3, cpr-1*, and *ugt-1* in excess zinc was significantly reduced in *hizr-1(lf)* and *hlh-30(lf)* mutants (Figure 4G). We further investigated the zinc-dependent *in vivo* activation mediated by *hizr-1* and *hlh-30* in the class HE gene *ugt-1*; we generated transgenic animals with the *ugt-1* promoter driving expression of nuclear localized mCherry. The transgene was introduced into wild type, *hizr-1(lf),* and *hlh-30(lf)* mutants. In wild-type animals, intestinal fluorescence derived from the *ugt-1* promoter was not detectable in replete zinc but robustly accumulated in intestinal nuclei in animals cultured in 200μM supplemental zinc (Figure 4D). By contrast, fluorescence in intestinal cells was not observed in *hizr-1(lf)* and *hlh-30(lf)* mutants cultured in excess zinc, indicating that both genes are necessary for full activation (Figure 4E-F). These results validate the existence of two classes of *hizr-1* target genes – class H are independent of *hlh-30*, whereas class HE are dependent on *hlh-30* (Fig 4H).

These results define a gene regulatory network controlled by HIZR-1 and HLH-30 constituted by 106 genes. These genes are significantly activated (*p*-value <0.001) in wild type animals exposed to excess zinc, compared to replete zinc conditions, measured by qRT-PCR. Figure 5 shows 12 class H genes that are predicted to be directly regulated by HIZR-1 but not HLH-30, including *cdf-2* and *ttm-1b*, and 94 class HE genes that are predicted to be directly regulated by HIZR-1 and HLH-30, including *sqst-3* and *ugt-1*. The class HE genes include several groups that are functionally related, including 18 genes that play a role in lysosome-autophagy biology, the most represented functional category. These results led us to hypothesize that excess zinc, acting through HIZR-1 and HLH-30, activates a set of genes that promote increased lysosome number and lysosome remodeling to accomplish high zinc homeostasis.

**Figure 5.**
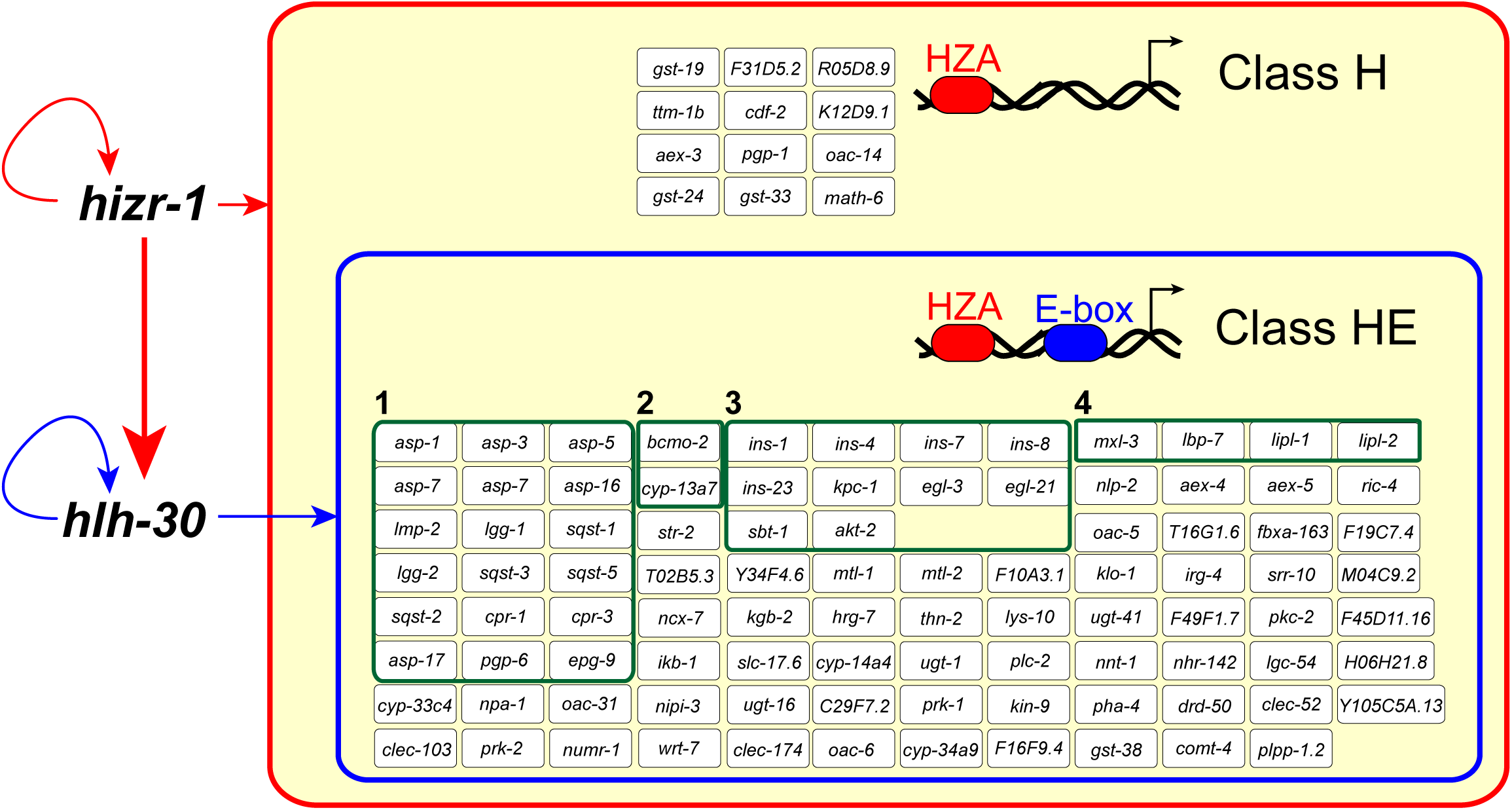
HIZR-1 and HLH-30 control a network of zinc-activated genes. A bioinformatic analysis of the promoters of zinc-activated genes identified in this study identified 106 with a match to the HZA enhancer (inside red square). We propose these genes are regulated by HIZR-1, which also regulates its own promoter and the promoter of *hlh-30* (red arrows). 94 genes were defined as class HE because the promoter contains a match to the E-box enhancer (inside blue rectangle). We propose these genes are regulated by HLH-30, which also regulates its own promoter (blue arrows). 12 genes were defined as class H because the promoter did not contain a match to the E-box enhancer (outside blue rectangle). Cartoons show DNA as a black helix, transcription start site as a black arrow, and enhancers as red (HZA) or blue (E-box) ovals. Green boxes indicate genes that have similar predicted functions based on gene ontology analysis: autophagy-lysosomal pathway (1), carotenoid metabolism (2), insulin signaling (3), and lipolysis (4).

### *hizr-1* and *hlh-30* determine remodeling of LROs and are necessary for the biogenesis of the expansion compartment in excess zinc

In replete zinc, *C. elegans* LROs can be detected in transgenic animals that express CDF-2 fused to GFP *(Pcdf-2::CDF-2::GFP)* from an array inserted in the genome *(amIs4)* by analyzing LysoTracker and CDF-2::GFP by fluorescence microscopy; in that condition, both markers are highly overlapping (Roh et al. 2012; Mendoza et al. 2024).

In excess zinc, LROs increase in number and display remodeling characterized by an increased volume of the expansion compartment – a structure named the bilobed granule (Roh et al. 2012; Shomer et al. 2019; Mendoza et al. 2024). The mechanisms necessary to form these bilobed granules are not completely understood. In excess zinc, the expansion compartment is labelled by CDF-2::GFP but not LysoTracker, which makes it possible to discriminate between the acidified and the expansion compartments; these compartments are separated by a region named the interface membrane. Our results showing that HIZR-1 and HLH-30 coordinately regulate multiple lysosome/autophagy genes suggests the model that these transcription factors mediate lysosome remodeling in response to excess zinc. To test this model, we used super-resolution microscopy to examine LROs in *hizr-1(lf)*, *hlh-30(lf)*, and *hlh-30(lf);hizr-1(lf)* double mutant animals carrying the *amIs4* inserted array. Synchronized L4 transgenic animals expressing CDF-2::GFP were shifted from zinc replete to zinc excess medium for ∼16 hours, stained with LysoTracker, and intestinal cells were imaged by fluorescence microscopy. In zinc replete medium, control animals displayed spherical LROs that were LysoTracker positive, indicating they are acidified; CDF-2::GFP was localized in a highly overlapping pattern, indicating the volume of the expansion compartment is minimal (Figure 6A). In excess zinc, the expansion compartment, which is CDF-2::GFP positive but LysoTracker negative, significantly increased in volume (Figure 6B). Almost every LRO that is LysoTracker positive has a high-volume expansion compartment; there are a small number of vesicles that are positive for CDF-2::GFP but negative for LysoTracker; we hypothesize these vesicles can fuse to the expansion compartment.

**Figure 6.**
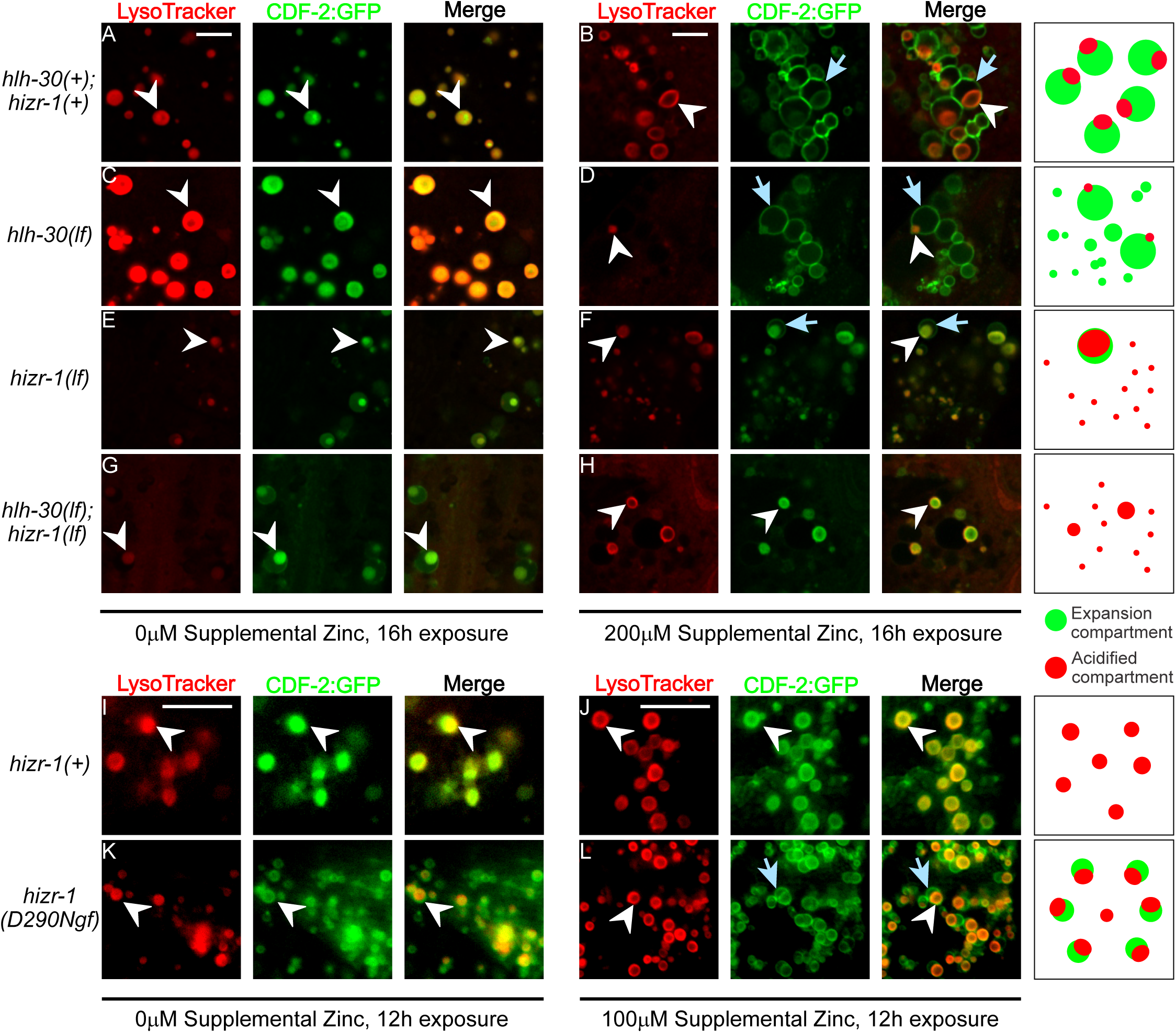
In excess zinc, HLH-30 promotes increased numbers of acidified compartments, whereas HIZR-1 promotes expansion compartments. (A-H) Animals cultured with 0μM supplemental zinc were shifted to 0 or 200μM supplemental zinc at the L4 stage and imaged by super-resolution microscopy after 16 hours. All strains contained *amIs4*, an integrated multicopy array that expresses CDF-2::GFP. Genotypes were *hizr-1(am286lf)*, *hlh-30(tm1978lf)*, and *hlh-30(tm1978lf); hizr-1(am286lf)*. Representative fluorescent images show LysoTracker (red), CDF-2::GFP (green), or merge (yellow overlap). Scale bar =5µm. In 0μM supplemental zinc, LROs in control animals appear to be spherical, and LysoTracker fluorescence (white arrowhead) and CDF-2::GFP fluorescence (white arrowhead) fully overlap (A,C,E,G). In 200μM supplemental zinc, LROs in control animals are remodeled and display a prominent expansion compartment (blue arrow) attached to the acidified compartment (white arrowhead) (B,D,F). Cartoons summarize the merge image, with red indicating acidified compartments and green indicating expansion compartments and expansion compartment vesicles. (I-L) Animals cultured with 0μM supplemental zinc were shifted to 0 or 100μM supplemental zinc at the L4 stage and imaged by standard epifluorescence microscopy after 12 hours. Genotypes were *hizr-1(+);amIs4* or *hizr-1(D290Ngf)*; *amIs4*. Representative fluorescent images show LysoTracker (red), CDF-2::GFP (green), or merge (yellow overlap). Scale bar = 5µm. In 100μM supplemental zinc in *hizr-1(+)* animals, LROs appear to be spherical, and LysoTracker fluorescence (white arrowhead) and CDF-2::GFP fluorescence (white arrowhead) fully overlap; in these conditions, *hizr-1(gf)* animals display an inflated expansion compartment (blue arrow). Representative images from at least two independent experiments per strain per condition.

The *hlh-30(lf);amIs4* mutant animals in zinc replete medium appeared similar to control; spherical LROs displayed both LysoTracker and CDF-2::GFP (Figure 6C). In zinc excess, *hlh-30(lf)* mutant animals displayed a reduced number of LysoTracker positive LROs, and the few that were detected displayed a small acidified compartment. In addition, these animals displayed a large number of CDF-2::GFP positive vesicles lacking acidified compartments (Figure 6D). These findings suggest that *hlh-30* is necessary to increase the number of acidified compartments in response to excess zinc. Furthermore, in the absence of a sufficient number of acidified compartments, the expansion compartment vesicles are more abundant and remain unfused, suggesting *hlh-30* promotes the fusion of the expansion compartment vesicles.

The *hizr-1(lf);amIs4* mutant animals in zinc replete medium appeared somewhat different from control; LROs displayed a small expansion compartment that is not typically observed in zinc replete conditions (Figure 6E). In zinc excess, *hizr-1(lf);amIs4* mutant animals displayed relatively small expansion compartments compared to control (Figure 6F). These finding suggest that *hizr-1* is necessary to increase the number and size of the expansion compartment vesicles in response to excess zinc. There seems to be a correlation between the number of vesicles available for fusion and the volume of the expansion compartments.

*hlh-30(lf);hizr-1(lf);amIs4* double mutant animals in zinc replete medium appeared similar to *hizr-1(lf)* single mutant animals; LROs displayed a small expansion compartment (Figure 6G). In zinc excess, *hlh-30(lf); hizr-1(lf)* double mutant animals displayed a strikingly abnormal pattern (Figure 6H). There were fewer LysoTracker positive LROs, similar to *hlh-30(lf)* mutants, and surprisingly there were no detectable expansion compartments. These findings suggest that *hlh-30* and *hizr-1* function cooperatively to mediate lysosome remodeling and bilobed granule biogenesis, since the double mutant animals display more severe defects that either of the single mutant animals and lack detectable expansion compartments.

To determine if *hizr-1* is sufficient to promote the increased volume of the expansion compartment, we analyzed the gain-of-function mutation *hizr-1(D290Ngf*). This allele contains a missense change in the ligand-binding domain of HIZR-1 which causes mutant protein to accumulate in the nucleus and activate target genes in zinc replete conditions (Warnhoff et al. 2017). In zinc replete conditions, *amIs4* animals displayed characteristic LROs that had highly overlapping staining with LysoTracker and CDF-2::GFP (Figure 6I). By contrast, the *hizr-1(D290Ngf)* mutant animals displayed LROs that had partially overlapping staining with LysoTracker and CDF-2::GFP; many LROs appeared to have a small expansion compartment, and these animals displayed a large number of vesicles that stained with CDF-2::GFP but not LysoTracker (Figure 6K). We interpret these CDF-2::GFP positive vesicles as expansion compartment vesicles that have not fused with LROs. Taking advantage of this sensitized genetic background, we used a moderate excess zinc condition (100μM) and a shorter response time (12 hours) to evaluate if the mutant activated HIZR-1 could trigger the appearance of bilobed granules. In these conditions, *amIs4* animals displayed spherical LROs with highly overlapping LysoTracker and CDF-2::GFP (Figure 6J), indicating that 100µM excess zinc is not a sufficient concentration to induce the appearance of bilobed granules. By contrast, *hizr-1(D290Ngf)* mutant animals displayed vesicles that stained with CDF-2::GFP but not LysoTracker, as observed in zinc replete medium, as well as LROs with expansion compartments of increased volume (Figure 6L). Thus, *hizr-1* is sufficient to promote the generation of expansion compartment vesicles in zinc replete conditions and induce/mediate the increase in the volume of the expansion compartment in moderate zinc excess.

## Discussion

### Genome-wide analysis of zinc regulated transcription reveals zinc influence on metabolism and signaling

Previous studies of high zinc-regulated transcription in *C. elegans* identified two key elements in the response pathway; the HZA enhancer present in the promoters of genes transcriptionally activated in excess zinc, and the HIZR-1 transcription factor that is regulated by zinc binding and interacts directly with the HZA enhancer. These discoveries were made using a small number of candidate genes that are activated by excess zinc, including *cdf-2*, *mtl-1*, *mtl-2,* and *ttm-1* (Roh et al. 2015; Warnhoff et al. 2017). However, the full extent of zinc-regulated transcription had not been defined. A comprehensive set of target genes is essential to understand all the dimensions of the organismal response to high zinc stress. Here we used two different strategies – RNA seq and microarrays – to define the genome-wide set of transcripts regulated by high zinc. We identified ∼1000 transcripts that display significantly higher levels and ∼520 transcripts that display significantly lower levels in response to excess zinc. This set included the established genes regulated by high zinc and many that have not been previously described. Transcriptional regulation by high zinc depends on several variables including the concentration of zinc, the duration of zinc exposure, the growth conditions, and the stage of the animals. The RNA seq experiment used 200μM zinc for 16-18 hours with mixed stage animals grown on solid medium with live *E. coli* as a food source. By contrast, the microarray experiment used 500μM zinc for at least six days to achieve steady state gene expression in defined liquid medium (CeMM), and fourth larval stage animals were synchronized by sorting prior to analysis. Both approaches identified the same set of strongly induced genes; these genes are likely to be induced in a wide range of excess zinc conditions. In addition, each experiment identified some unique genes whose activation or repression by excess zinc may be condition dependent.

While there is an emerging understanding of the mechanisms that promote activation of gene expression in high zinc, the mechanisms that cause repression in high zinc are not established. We previously identified one gene that is repressed in high zinc: *zipt-2.3* encodes a transporter that localizes to lysosome-related organelles in the intestine and releases stored zinc to the cytosol (Mendoza et al. 2024). The identification of ∼520 additional genes that are repressed by high zinc is an important step in deciphering the mechanisms, since these genes may share common regulatory control. The regulation of these genes might occur by decreasing transcription initiation or increasing transcript degradation, and these discoveries lay the foundation for future experiments to determine the mechanism of regulation.

The genes that are activated by excess zinc represent multiple functional categories, and here we highlight four functional categories: autophagy-lysosome pathway, carotenoid metabolism, insulin signaling, and lipolysis. Because lysosome-related organelles in intestinal cells are a site of zinc storage in *C. elegans*, we focused on the function of the autophagy-lysosome pathway.

### HLH-30/TFEB, the master regulator of lysosome biogenesis, is directly activated by HIZR-1 and mediates high zinc homeostasis

Transcription factor EB (TFEB) was identified in mammals as the master regulator of autophagy and lysosome biogenesis (Settembre et al. 2011; Roczniak-Ferguson et al. 2012; Lapierre et al. 2013; Settembre et al. 2013). This helix-loop-helix transcription factor controls multiple genes that encode components of lysosomes. While it is possible to form acidified lysosomes in TFEB mutants, the rapid expansion of lysosome numbers in stressful conditions such as starvation is not possible in TFEB mutants (Settembre et al. 2013). *C. elegans* contains a single homolog of TFEB, called HLH-30 (Lapierre et al. 2013). *hlh-30* mutants display a wide range of phenotypes consistent with roles in lysosome biogenesis and autophagy, such as hypersensitivity to starvation (Settembre et al. 2013). However, *hlh-30* has not been previously implicated in high zinc homeostasis. Here we show that *hlh-30* transcripts accumulate significantly in high zinc. The *hlh-30* promoter contains a predicted HZA enhancer, and activation of *hlh-30* in high zinc requires *hizr-1*. Thus, *hlh-30* appears to be a direct target of *hizr-1* that is significantly activated in high zinc. To monitor the function of HLH-30, we analyzed nuclear accumulation of an HLH-30::RFP protein. HLH-30 accumulated in intestinal nuclei, an indication of functional activation by this transcription factor. The regulation of TFEB has been analyzed in mammals, and phosphorylation that mediates nuclear localization has been suggested to be an important control point (Li et al. 2018; Trivedi et al. 2020; Zhang et al. 2023). Here we show that a different mechanism, transcriptional regulation, appears to be an important regulatory control point for HLH-30.

To determine the function of *hlh-30* during high zinc homeostasis, we analyzed a strong loss-of-function mutation. *hlh-30(lf)* mutants were hypersensitive to excess zinc, displaying a significant growth defect. Thus, *hlh-30* is necessary for these animals to tolerate high zinc stress. We previously demonstrated that *hizr-1(lf)* mutants display a similar growth defect in excess zinc. Thus, both genes play important and independent roles during high zinc homeostasis. To elucidate the interaction between these two genes, we generated and analyzed double mutants. These animals were extremely hypersensitive to excess zinc and displayed growth defects under conditions where both single mutants were relatively normal. These results indicate that *hlh-30* and *hizr-1* also play redundant roles in high zinc tolerance.

Since HIZR-1 and HLH-30 are both transcription factors responsive to excess zinc whose DNA binding motifs are known, we investigated the possibility that some target genes are coordinately regulated by both. We searched for genes that are activated by excess zinc and have an HZA enhancer and an E-box enhancer, which we named class HE target genes. We identified many that fit these criteria, including 18 genes likely involved in lysosome biogenesis. In addition, we identified class H target genes that have an HZA enhancer but not an E-box enhancer. To validate regulatory control, we analyzed transcription in wild type, *hizr-1(lf)* mutants, and *hlh-30(lf)* mutants. Three class H genes, *cdf-2*, *asp-17*, and *cyp-14a4* required *hizr-1* but not *hlh-30* for full activation by excess zinc. By contrast, three class HE genes, *sqst-3*, *cpr-1*, and *ugt-1*, required both *hizr-1* and *hlh-30* for full activation by excess zinc. These results establish a transcriptional network that controls gene expression in response to excess zinc. HIZR-1 function as the high zinc sensor, directly binding zinc under excess zinc conditions and translocating to the nucleus to activate genes controlled by an HZA enhancer. One such gene is HLH-30, and activation of HLH-30 results in increased transcription of genes that contain the E-box enhancer. Based on this model, we propose HIZR-1 and HLH-30 coordinately regulate class HE genes, whereas HIZR-1 but not HLH-30 regulates class H genes. We speculate that HLH-30 but not HIZR-1 regulates genes with only the E-box enhancer. Thus, excess zinc initiates a transcription factor cascade to control multiple target genes.

### *hizr-1* and *hlh-30* mediate remodeling of lysosome-related organelles

Because zinc is an essential element, organisms evolved mechanisms to store excess zinc during times of plenty so that it can be mobilized during times of deficiency. Lysosomes are emerging as an evolutionary conserved site for storage of excess zinc, including the zincosome in mammals, the vacuole in yeast, the acidocalcisome in Chlamydomonas, and the gut granules of *C. elegans* (Zalewski et al. 1993; Coyle et al. 1994; MacDiarmid et al. 2000; MacDiarmid et al. 2003; Simm et al. 2007; Ferella et al. 2008; Wellenreuther et al. 2009; Yagisawa et al. 2009; Roh et al. 2012; Mendoza et al. 2024). We previously showed that *cdf-2*, a class H target gene that is directly regulated by HIZR-1, encodes a zinc transporter that stores zinc in lysosome-related organelles in intestinal cells (Davis et al. 2009; Roh et al. 2012; Roh et al. 2015). In addition to increased levels of CDF-2, LROs are remodeled in excess zinc with the appearance of a prominent expansion compartment. Based on super-resolution microscopy studies, Mendoza et al. (2024) proposed that the expansion compartment increases in volume by fusion of vesicles containing CDF-2 (Mendoza et al. 2024). The role of the expansion compartment is to facilitate a rapid shift in the ratio of zinc transporters on the membrane of LROs. However, the mechanisms that mediate LRO remodeling are only beginning to be defined. Here we used CDF-2 to mark the expansion compartment and LysoTracker to mark the acidified compartment. We showed that *hizr-1(lf)* mutants display a defect in increasing the volume of the expansion compartment, indicating that *hizr-1* is necessary for this process. Furthermore, we showed that a *hizr-1(gf)* mutation is sufficient to increase the volume of the expansion compartment in conditions where this does not occur in wild-type animals. In *hlh-30* mutants, vesicles that are positive for CDF-2 accumulate, and this accumulation is abrogated in *hizr-1(lf)* mutant animals. We propose that *hizr-1*, acting through class H target genes, mediates the formation of CDF-2 positive vesicles that fuse with the expansion compartment and increase its volume during zinc excess (Figure 7). Thus, *hizr-1* not only activates transcription of the *cdf-2* gene, it also promotes the formation of CDF-2 positive vesicles that fuse with the expansion compartment to deliver CDF-2 transporter and increase the capacity for zinc storage. The *hizr-1* target genes that mediate the formation of these vesicles remain to be determined and are an important objective of future research.

**Figure 7.**
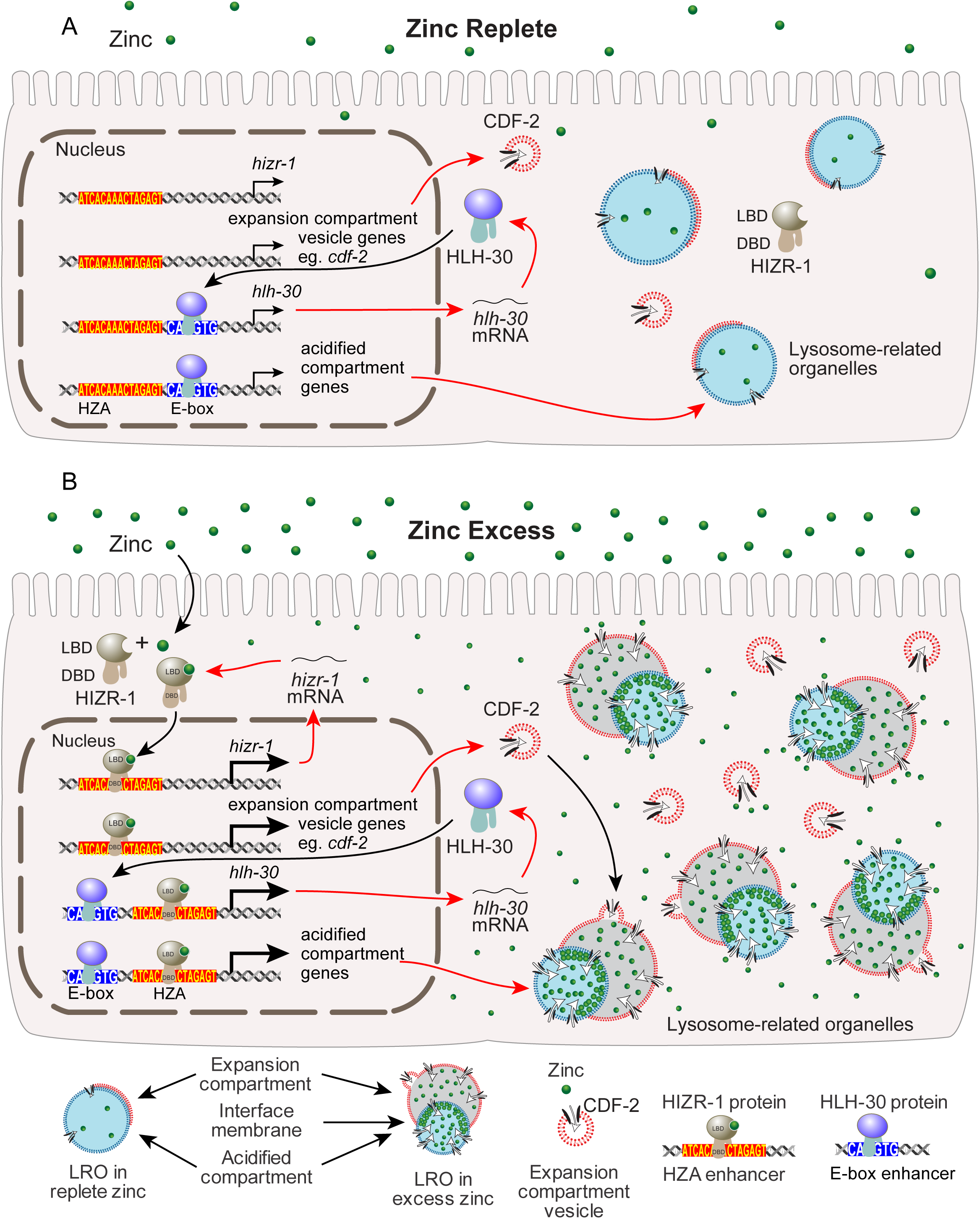
A model of HIZR-1 and HLH-30 regulating lysosome remodeling in excess zinc. A) In zinc replete conditions, HIZR-1 and HLH-30 proteins are primarily localized in the cytoplasm. A low level of nuclear localized proteins mediates basal transcription; HIZR-1 binds the HZA enhancer, whereas HLH-30 binds the E-box enhancer. Lysosomes are composed of prominent acidified compartments (blue) and small volume expansion compartments (red); there are a small number of CDF-2 positive expansion compartment vesicles (red with black CDF-2 transporter – white arrow indicates the direction of zinc transport by CDF-2). B) In excess zinc conditions, zinc accumulates in the cytoplasm of the intestinal cells and binds the ligand binding domain (LBD) of HIZR-1, triggering nuclear localization, binding to the HZA enhancer through the DNA-binding domain (DBD), and transcriptional activation of target genes. Target genes include class H and class HE genes, including *hizr-1*, *cdf-2*, and *hlh-30*. HIZR-1 target genes promote the formation of CDF-2 positive vesicles that fuse to the expansion compartment, thereby increasing its volume. The *hlh-30* gene is activated by *hizr-1* in excess zinc, and HLH-30 protein translocates to the nucleus and binds the E-box enhancer. Target genes include class HE genes, such as *hlh-30* which autoregulates. HLH-30 target genes promote formation of acidified compartments. The combined action of HIZR-1 and HLH-30 promotes the formation of lysosomes with high volume expansion compartments.

*hlh-30(lf)* mutants display a different defect in LRO remodeling. In excess zinc, *hlh-30(lf)* mutants have a reduced number of acidified compartments that stain with LysoTracker. Thus, while *hlh-30(lf)* is not necessary to generate some acidified compartments, *hlh-30* is necessary to increase the number of acidified compartments in response to high zinc stress. This is consistent with the role of TFEB in controlling lysosome number in response to stress situations such as starvation. The other striking change in *hlh-30(lf)* mutants is the accumulation of CDF-2 positive vesicles that are not fused to acidified compartments, a phenomenon also observed in worms treated with *pgp-2* and *glo-3* RNAi (Roh et al. 2012). We speculate this reflects an imbalance between the small number of acidified compartments and the large number of vesicles. It is also possible that *hlh-30* is necessary for efficient fusion of vesicles to the acidified compartment, which might reflect a role for *hlh-30* target genes that contain only an E box and drive vesicle fusion, such as snare proteins. These results provide fresh insights into the genetic control of LRO remodeling. In excess zinc, LROs increase in number and are remodeled to have a high-volume expansion compartment. *hlh-30* mediates the increase in the number of acidified compartments, whereas *hizr-1* mediates the increase in the number of vesicles that fuse with the expansion compartment to increase its volume. In *hlh-30* mutants, there are fewer LROs and an excess of CDF-2 positive vesicles, resulting in reduced zinc storage and hypersensitivity to excess zinc. In *hizr-1(lf)* mutants, the expansion compartment does not increase in volume, suggesting a failure to deliver CDF-2 transporter. This defect compromises zinc storage, resulting in hypersensitivity to excess zinc. Thus, the coordinated action of *hlh-30* and *hizr-1* is necessary for LRO remodeling and high zinc tolerance.

## Experimental procedures

### General methods and strains

*C. elegans* strains were cultured at 20°C on nematode growth medium (NGM) seeded with *Escherichia coli* OP50 except as noted. The wild-type *C. elegans* and parent of all mutant strains was Bristol N2. We used the following chromosomal mutations: *hizr-1(am286lf)*, *hizr-1(am285gf)* (Warnhoff et al. 2017), and *hlh-30(tm1978lf)* (Murphy et al. 2019); and previously described transgenic strains: *amIs4* (*cdf-2::gfp + unc-119*(+)) (Roh et al. 2012), HT1704 (*ins-8p::gfp + unc-119*(+)) (Ritter et al. 2013), *stIs11634* (*ugt-1p::H1-mCherry + unc-119*(+)), VL749 *wwIs24 [Pacdh1::GFP + unc-119(+)]* (MacNeil et al. 2013), *hlh-30p::hlh-30::rfp* in *hlh-30(tm1978lf)* (Murphy et al. 2019), and *hizr-1p::gfp* (Warnhoff et al. 2017). Double mutant animals were generated by standard methods, and genotypes were confirmed by PCR or DNA sequencing. Mutants were obtained from the *Caenorhabditis* Genetics Center (CGC) and the National Bioresource Project. The full list of the strains used is provided in Table S14.

### Plasmid DNA construction and transgenic strain generation

To generate transcriptional fusion constructs for *bcmo-2* (P*bcmo-2*::*gfp*), *thn-2* (P*thn-2::gfp*), and *cyp-14a4* (P*cyp-14a4*::*gfp*), we polymerase chain reaction (PCR)-amplified DNA fragments positioned upstream of the coding sequence using wild-type *C. elegans* DNA. The *bcmo-2* promoter was amplified from the ATG start codon to 1,371 base pairs upstream of the ATG start codon. The *thn-2* promoter was amplified from the ATG start codon to 717 base pairs upstream of the ATG start codon. The *cyp-14a4* promoter was amplified from the ATG start codon to 642 base pairs upstream of the ATG start codon. The oligonucleotide primers are described in Table S15. These fragments were ligated into pBluescript SK+ (Stratagene) containing the green fluorescent protein (GFP) coding sequence and the *unc-54* 3’ untranslated region (UTR). The resulting plasmids pCAC1 (P*bcmo-2*::*gfp*), pCAC2 (P*thn-2::gfp),* and pCAC3 (P*cyp-14a4*::*gfp*) were confirmed by DNA sequencing. Transgenic animals containing extrachromosomal arrays were generated by injecting the gonad of worms with a plasmid of interest and the co-injection marker plasmid pRF4 which encodes the dominant ROL-6(R71C GF) mutant protein (Mello et al. 1991). Transgenic animals were selected by the Rol phenotype.

### Analysis of gene expression levels by RNA sequencing using WT and *hizr-1(lf)* animals cultured on NAMM dishes with replete and excess zinc

To prepare RNA, we cultured 3 wild type L4 stage hermaphrodites at 20°C on an NGM dish seeded with 1X *E. coli* OP50 – 40 dishes were prepared. In parallel, we cultured 3 *C. elegans hizr-1(am286lf)* L4 stage hermaphrodites on an NGM dish seeded with 1X *E. coli* OP50 – 40 dishes were prepared. After 4 days, the mixed-stage populations were collected by washing with M9 buffer and cultured on the same number of NAMM dishes seeded with 10X *E. coli* OP50 supplemented with either 0μM ZnSO_4_ (40 dishes) or with 200μM ZnSO_4_ (40 dishes) as follows: 20 dishes of wild type with 200μM ZnSO_4_, 20 dishes of wild type without supplemental ZnSO_4_, 20 dishes of *hizr-1(lf)* with 200μM ZnSO_4_, and 20 dishes of *hizr-1(lf)* without supplemental ZnSO_4_. Animals were cultured for 16-18h at 20°C and collected by washing with M9; animals from 10 dishes were pooled into one biological replicate. This procedure generated two biological replicates for each wild type and *hizr-1* mutant condition, for a total of 8 samples. RNA was isolated using TriZol (Invitrogen) as previously described (Warnhoff et al. 2017) with minor modifications, including additional freeze-thaw cycles and additional chloroform extractions to reduce protein contamination. RNA from each sample was treated with DNaseI (Fermentas), purified using an RNeasy kit (Invitrogen), and eluted in 20μl RNase/DNase free water. RNA quality of each sample was estimated by chromatography, and all samples had a RNA integrity number (RIN) value of > 10.

Library preparation was performed with 1μg of total RNA; RNA concentration was determined by Qubit, and integrity was determined using an Agilent tapestation or bioanalyzer. Ribosomal RNA was removed by a hybridization method using Ribo-ZERO kits (Illumina). Depletion and mRNA yield was confirmed by bioanalyzer. mRNA was then fragmented in buffer containing 40mM Tris Acetate pH 8.2, 100mM Potassium Acetate and 30mM Magnesium Acetate and heating to 94 degrees for 150 seconds. The mRNA was reverse transcribed to yield cDNA using SuperScript III RT enzyme (Life Technologies, per manufacturer’s instructions) and random hexamers. A second strand reaction was performed to yield ds-cDNA. The cDNA was blunt ended, an A base was added to the 3’ ends, and Illumina sequencing adapters were ligated to the ends. Ligated fragments were amplified for 12-15 cycles using primers incorporating unique index tags. Library molarity was determined by Qubit assay for concentration and tapestation for size. An equimolar pool was made of all libraries with unique indices. Fragments were sequenced on an Illumina HiSeq-3000 using single reads extending 50 bases.

Basecalls and demultiplexing were performed with Illumina’s bcl2fastq software and a custom python demultiplexing program with a maximum of one mismatch in the indexing read. RNA-seq reads were then aligned to the Ensembl release 76 top-level assembly with STAR version 2.0.4b (Dobin et al. 2013). Gene counts were derived from the number of uniquely aligned unambiguous reads by Subread:featureCount version 1.4.5 (Liao et al. 2014). Sequencing performance was assessed for the total number of aligned reads, total number of uniquely aligned reads, and features detected. The ribosomal fraction, known junction saturation, and read distribution over known gene models were quantified with RSeQC version 2.3 (Wang et al. 2012).

All gene counts were imported into the R/Bioconductor package EdgeR (Robinson et al. 2010), and TMM normalization size factors were calculated to adjust for samples for differences in library size. Genes or transcripts not expressed in any sample were excluded from further analysis. The TMM size factors and the matrix of counts were imported into the R/Bioconductor package Limma (Ritchie et al. 2015). Weighted likelihoods based on the observed mean-variance relationship were derived for every gene with Voom (Liu et al. 2015). The performance of all genes was assessed with plots of the residual standard deviation of every gene to their average log-count with a robustly fitted trend line of the residuals. Differential expression analysis was performed to analyze for differences between conditions, and the results were filtered for only those genes with Benjamini-Hochberg false-discovery rate adjusted *p*-values less than or equal to 0.05.

### Analysis of gene expression levels by microarray using wild type animals cultured in CeMM medium with replete and excess zinc

Large populations of wild type worms were cultured in fully defined *C. elegans* maintenance medium (CeMM, (Davis et al. 2009)) containing 30μM zinc chloride (ZnCl_2_) for multiple generations to achieve steady state levels of gene expression. Worms were collected by centrifugation and washed 2X in M9 buffer/CeMM without ZnCl_2_. The pelleted worms were transferred to 5ml CeMM containing 30μM ZnCl_2_ in a 25cm^2^ T-flask, or to 15 ml CeMM containing 500μM ZnCl_2_ in a 75cm^2^ T-flasks. Worms were cultured for 6 days at 20°C to achieve steady state levels of gene expression. In these culture conditions, 30μM ZnCl_2_ is replete whereas 500μM ZnCl_2_ is excess (Davis et al. 2009).

To obtain populations of worms synchronized at the L4 stage, we diluted mixed stage populations of worms in 50mM Tris HCl containing the corresponding 30 or 500μM ZnCl_2_ and sorted by time of flight using a COPAS-BIOSORT. Approximately 4000 L4 worms were isolated from independent cultures and pooled to generate a single biological replicate; we generated 4 biological replicates each for the 30μM and 500μM ZnCl_2_ populations.

Total RNA from the L4 stage worms was isolated using TRIzol (Invitrogen) and treated with DNAse I (Turbo DNA Free kit, Applied Biosystems). RNA was quantified by measuring absorbance at 260nm and 280nm and by using the Agilent 2100 bioanalyzer (Agilent technologies).

RNA transcripts were first amplified by T7 linear amplification (MessageAmp aRNA amplification kit; Ambion). 1μg of RNA was primed with an oligo-dT T7 primer (1μl) by heating to 70°C for 10min and cooling on ice for 3min. For 1^st^ strand cDNA synthesis, each RNA received 10X reaction buffer (2μl), dNTP mix (4μl), RNase inhibitor (1μl) and reverse transcriptase (1μl) (Superscript II; Invitrogen). Reverse transcription was carried out for 2hours at 42°C. After a 3min incubation on ice, the cDNA underwent 2^nd^ strand synthesis by adding water (63μl), 10X 2^nd^ strand buffer (10μl), dNTP mix (4μl), DNA polymerase (2μl), and RNAse H (1μl). This mixture was incubated at 16°C for 2h. Following a short column cleanup (DNA clean & concentrator-5, Zymo Research), *in vitro*-transcription was carried out by adding non-modified bases (ATP, CTP, GTP, and UTP, 4μl each), 10X T7 reaction buffer (4μl), and T7 RNA polymerase enzyme mix (4μl). The reaction was carried out for 14h. Following reaction termination with water (60μl), the aRNAs were cleaned with RNeasy columns (Qiagen). The aRNAs were chemically labeled with Kreatech ULS RNA labeling kit (Kreatech Diagnostics); 3μg of each RNA dissolved in water was mixed with the Kreatech 10X labeling buffer (2μl) and Kreatech cy/DY-ULS (either cy3 or cy5) (2μl). The reactions were incubated at 85°C for 15min in the dark and placed on ice for 3min. Labeled aRNAs were purified with Kreapure gel columns and fragmented (Fragmentation reagents; Ambion). The labeled RNAs were then quantitated on a NanoDrop spectrophotometer.

For hybridization, 2μg of each labeled/fragmented aRNA was paired then suspended in formamide-based hybridization buffer (vial 7-Genisphere) (26μl). The hybridization solution was heated to 70°C for 5min, allowed to cool to room temperature, and added to the microarray under a supported glass coverslip (Erie Scientific) at 43°C for 16-20h at high humidity. Slides were gently submerged into 2X SSC, 0.2% SDS (at 43°C) for 11min, transferred to 0.2X SSC (at room temperature) for 11min, and spin dried by centrifugation. To prevent fluorophore degradation, the arrays were treated with Dyesaver (Genisphere).

Slides were scanned on a Perkin Elmer ScanArray Express HT scanner to detect Cy3 and Cy5 fluorescence. Laser power was kept constant for Cy3/Cy5 scans and PMT was varied for each experiment based on optimal signal intensity with lowest possible background fluorescence. A low pmt setting scan was also performed to recover signal from saturated elements. Gridding and analysis of images was performed using ScanArray v3.0 (Perkin Elmer).

### Determination of gene expression levels by quantitative real-time reverse transcription polymerase chain reaction (qRT-PCR) using animals cultured on NAMM dishes in replete and excess zinc

RNA used for qRT-PCR was purified as described above for RNA-sequencing. RNA was prepared from populations of wild type, *hizr-1(lf)* mutants, and *hlh-30(lf)* mutants. Animals were cultured on standard NGM dishes, collected by washing, and cultured at 20°C on NAMM dishes seeded with 10X concentrated *E. coli* OP50 and supplemented with 0μM and 200μM ZnSO_4_ for 16-24h.

For cDNA synthesis, we used 0.5 - 1μg of total RNA with the High-capacity cDNA Reverse Transcription kit according to the manufacturer’s protocol. The synthesized cDNA was diluted 1:2, and 2μl of the diluted cDNA was used for each qRT-PCR reaction. qRT-PCR was performed using SYBR Green PCR Master Mix (Bio-rad) and an Applied Biosystems 7900 thermocycler as previously reported (Roh et al. 2015). Fold change was determined by comparing levels of target gene expression between different zinc conditions or between different genotypes. The reference gene was *ama-1* or *rps-23*. The fold change in gene expression was calculated using the ΔΔCT method as previously reported (Livak and Schmittgen 2001; Schmittgen and Livak 2008). Data CT values for the qRT-PCR experiments reported are included in are included in Tables S6, S7, S8, S10, S11, and S13. Oligonucleotide primers are listed in Table S15. Statistical analyses were performed in GraphPad Prism 10.

### Measuring animal growth rate in replete and excess zinc

Gravid adult hermaphrodites were treated with NaOH and bleach, and eggs were incubated in M9 solution overnight to allow hatching and synchronized arrest at the L1 stage. L1 stage animals were transferred to NAMM dishes supplemented with ZnSO_4_. After 3 days, animals were washed twice in M9, paralyzed with 25mM tetramisole hydrochloride in M9, and mounted on a 2% agarose pad on a microscope slide. Images were captured with a Zeiss 2 microscope equipped with a Zeiss AxioCam MRm digital camera. Length of individual animals was measured as a quantitative measure of growth using ImageJ software (NIH) by drawing a line from the tip of the nose to the tip of the tail.

### Confocal and super resolution microscopy of LROs

Lysotracker RED DND-99 (1mM, Invitrogen L7528) was diluted in 10X concentrated *E. coli* OP50 to a final concentration of 10μM and dispensed on NAMM dishes. L4 stage hermaphrodites were cultured on these dishes for 16-18 h in the dark, washed twice in M9, paralyzed with 25mM tetramisole hydrochloride in M9, and mounted on a 2% agarose pad on a microscope slide. Fluorescence was recorded by using a Zeiss Axioplan 2 microscope equipped with a Zeiss AxioCam MRm digital camera using identical settings and exposure times.

Live-worm super-resolution imaging was performed using the Zeiss LSM880 II with AiryScan. LROs were imaged using the 60x objective in a z-stack. CDF-2::GFP was detected using the 488 nm laser. LysoTracker Red was detected using the 561 nm laser. Following capture, images were deconvolved using AiryScan processing which permitted a resolution of ∼120 nm.

### Identification of HZA and E-box motifs in promoters of zinc-regulated genes

The HZA was initially defined based on the promoter analysis of four *C. elegans* genes, *mtl-1, mtl-2, cdf-2* and *ttm-1,* and their orthologs in 3 species. The HZA was refined and improved by subsequent analysis of 29 *C. elegans* Cd-induced genes which included these 4 genes (Roh et al. 2015). The E-box matrix for HLH-30 was obtained from the Cis-BP database (Weirauch et al. 2014). We used the Regulatory Sequence Analysis Tools (RSAT) website to obtain the predicted promoter sequence of zinc-regulated *C. elegans* genes that were identified by whole genome analysis and validated by qRT-PCR. The predicted promoter sequence is defined as the DNA region from the start codon upstream to the closest gene or −2000 bp. This list of DNA sequences was input into Cis-regulatory enriched regions (CRERs) to identify HZA and E-box motifs along the 2000bp upstream sequence and to search for modules located within a 50bp window (Turatsinze et al. 2008).

### Analysis of HLH-30 protein localization

Gravid adults containing the *hlh-30p::hlh-30::rfp* multicopy extrachromosomal array (*amEx272*) were bleached, and the eggs were directly dropped on NGM dishes with 1X *E. coli* OP50, to prevent starvation-induced expression of HLH-30::RFP. Worms were incubated for 48 h (approximately the L4 stage) at 20 °C, and ∼100 synchronized worms were moved onto NAMM dishes supplemented with 0µM or 200µM zinc. Animals were incubated for 16-18 h, worms were collected by picking, paralyzed with 25mM tetramisole hydrochloride in M9, and mounted on a 2% agarose pad on a microscope slide. Fluorescence was recorded by using a Zeiss Axioplan 2 microscope equipped with a Zeiss AxioCam MRm digital camera using identical settings and exposure times. Image analysis was done by using ImageJ; brightness and contrast were treated equally for zinc replete and zinc excess exposed worms. To estimate the percentage of worms having positive HLH-30::RFP nuclear accumulation after the incubation period (16-18 h) on NAMM dishes, we used stereoscopic microscopy to score the number of worms displaying RFP positive nuclei foci throughout the intestine versus the total number of worms.

### Analysis of localization of mammalian TFEB

HEK cells were cultured in Dulbecco’s Modified Eagle Medium (DMEM; Gibco, 10965084) supplemented with 10% fetal bovine serum (FBS) (Thermo Scientific 26140029), 1 mg/mL sodium pyruvate (Gibco, 1360070), 100 U/mL penicillin, and 100 μg/mL streptomycin (Thermo Scientific 15140122). Cells were seeded onto 4-well Lab-Tek chamber slides (Thermo Scientific, 177437) and transfected at approximately 50% confluence with 100 ng of pAcGFP1-mTFEB plasmid using Lipofectamine™ 3000 (Thermo Scientific, L3000015) according to the manufacturer’s protocol. Twenty hours post-transfection, the culture medium was replaced with either normal medium (Zinc Replete) or medium supplemented with 200 μM ZnSO (Zinc Excess). After 4 hours of incubation, cells were fixed with 4% paraformaldehyde and counterstained with DAPI. Fluorescence images were acquired using a ZEISS LSM 700 confocal microscope. Nuclear and total GFP fluorescence intensities were measured using ImageJ software.

### Analysis of transgenic animals with zinc-regulated promoters fused to GFP

Gravid adults for each strain were bleached, the eggs were dropped directly onto NGM dishes lacking *E. coli* OP50, and animals were allowed to starve for 18h to synchronize to L1 stage. Then they were transferred to NGM dishes with 1X *E. coli* OP50. L1 animals were incubated at 20°C on NGM dishes until they reached the L4 stage. Around 100 synchronized L4 worms carrying the corresponding markers (i.e. *rol-6* for extrachromosomal arrays) were collected by picking and moved to NAMM dishes supplemented with 0µM zinc or 200µM zinc, incubated for 16-18 h, paralyzed with 25mM tetramisole hydrochloride in M9, and mounted on a 2% agarose pad on a microscope slide. Fluorescence was recorded by using a Zeiss Axioplan 2 microscope equipped with a Zeiss AxioCam MRm digital camera using identical settings and exposure times. Image analysis was done by using ImageJ; brightness and contrast were treated equally for zinc replete and zinc excess exposed worms.

## Data Availability statement

The genomic data described in this publication have been deposited in NCBI’s Gene Expression Omnibus (Edgar 2002) and are accessible through GEO Series accession number GSE172079.

## Acknowledgements

This work was supported in part by the National Institutes of Health (R01 GM068598 to K.K.; 1K99GM146016-01 to A.D.M) and by CONACyT, Mexico (postdoctoral fellowship 238727 to C.C.). We thank the *Caenorhabditis* Genetics Center for providing strains and the Genome Technology Access Center in the Department of Genetics at Washington University School of Medicine for help with genomic analysis. We are grateful to Seth Crosby for support with microarray analysis and Peter Bayguinov, Clifford Luke, Stephen Pak, and Gary Silverman for microscopy and image analysis. This publication is solely the responsibility of the authors and does not necessarily represent the official view of CONACyT or NIH.

---

**Supplemental Figure 1 (with main Figure 1). Identification of zinc-regulated genes by microarray analysis.** Populations of WT worms were cultured in fully defined CeMM medium containing 30μM ZnCl_2_ for several weeks (Davis et al. 2009). To analyze transcription in zinc excess conditions, worms were transferred to CeMM containing 500μM ZnCl_2_ and cultured for 6 days. A synchronous population of L4 staged worms was obtained using a COPAS-BIOSORT. Four biological replicates were performed for 30μM and 500μM ZnCl_2_ CeMM. RNA was isolated and analyzed using microarrays containing ∼19,000 *C. elegans* genes. Each point is data for one gene, showing Log_2_ fold change of expression level and the P value. Blue and gray indicate significant and nonsignificant P values, respectively. Red indicates genes previously established to be zinc-activated.

**Supplemental Figure 2 (with main Figure 1). Comparison of zinc-regulated genes identified by RNA seq and microarray analysis.** (A,B) Venn diagrams show the overlap (red) between lists of genes identified by RNA seq (blue circle) and microarray (white circle). Panel A displays genes activated by excess zinc, and panel B displays genes repressed by excess zinc. For activated genes, ∼35% of the 130 genes identified by microarray were also identified by RNA seq. For repressed genes, ∼4% of the 113 genes identified by microarray were also identified by RNA seq. (C) A list of the overlapping genes with the Log_2_ Fold Change and p-value for the method of RNA seq and microarray. Fourteen of these activated genes were analyzed by qRT-PCR, and all displayed activation by this method.

**Supplemental Figure 3 (with main Figure 1). Excess manganese did not cause the same changes in gene expression as excess zinc.** Mixed-stage populations of WT worms were cultured on standard (replete for zinc and manganese), zinc excess (200μM supplemental zinc), or manganese excess (500μM supplemental manganese) NAMM dishes for 16-18 hours, and gene expression was analyzed by qRT-PCR. Bars indicate the ratio of normalized mRNA levels in zinc excess/zinc replete conditions (black) or in manganese excess/manganese replete conditions (white). Bars indicate Log_2_ fold change. A value of 0 indicates no change between conditions – positive values indicate activation, and negative values indicate repression. Statistical analysis is done on at least three independent experiments, using one way ANOVA with post-hoc Dunnett T3. For significant p values: *<0.05; **<0.01; ***<0.001; ****<0.0001. Non-significant p values are indicated as “ns”. Error bars represent mean ± SD Gene names are indicated below.

**Supplemental Figure 4 (with main Figure 1). Localization of four zinc activated genes *in vivo*.** Transgenic animals containing plasmids with the promoter of *cyp-14a4* (A-D), *thn-2* (E-H), or *ins-8* (M-P) driving GFP and *ugt-1* (I-L) driving H1::mCherry were cultured on zinc replete (0μM supplemental ZnSO_4_) or zinc excess (200μM supplemental ZnSO_4_) NAMM for 16-18 hours and visualized by bright field (upper) or fluorescence (lower) microscopy. Scale bar = 100µm. The *ins-8* strain includes *unc-119(+)* as a marker. Yellow arrowheads indicate fluorescent protein expression in zinc excess conditions. The results indicate that the promoters of *cyp-14a4*, *thn-2*, and *ugt-*1are not strongly expressed in zinc replete conditions and are activated specifically in intestinal cells by excess zinc. The promoter of *ins-8* displays expression in the anterior and posterior regions in zinc replete conditions, and it appears to be activated in the anterior intestine by excess zinc.

**Supplemental Figure 5 (with main Figure 4). *hlh-30* is not necessary for activation of the *hizr-1* promoter in excess zinc.** (A) The alignment shows DNA sequences from the promoters of *hlh-30*, *hizr-1*, *lipl-1*, *lipl-2, lipl-3,* and *sqst-3*. Gray indicates nucleotides identical in all six sequences. Position indicates the number of base pairs upstream of the translation start site. The *p*-value is defined as the probability of observing a match score as good when the motif is compared to a random sequence (Bailey and Gribskov 1998), a low *p*-value suggests the result is unlikely to be random. (B-I) Transgenic animals containing plasmids with the *hizr-1* promoter driving GFP were cultured on NAMM dishes with 0 or 100μM supplemental zinc for 6 hours and visualized by bright field (upper) or fluorescence (lower) microscopy. Scale bar = 100µm. *hizr-1* was activated by zinc excess in control (compare C to E) and *hlh-30(tm1978lf*) mutants (compare G to I). Although the *hizr-1* promoter contains a predicted E-box sequence, *hlh-30* is not necessary for activation of *hizr-1* transcription in excess zinc.

