## Supplementary Figures 1-5 for "Zinc excess promotes lysosome remodeling by activating HLH-30/TFEB through the action of the high zinc sensor HIZR-1"

### Figure S1

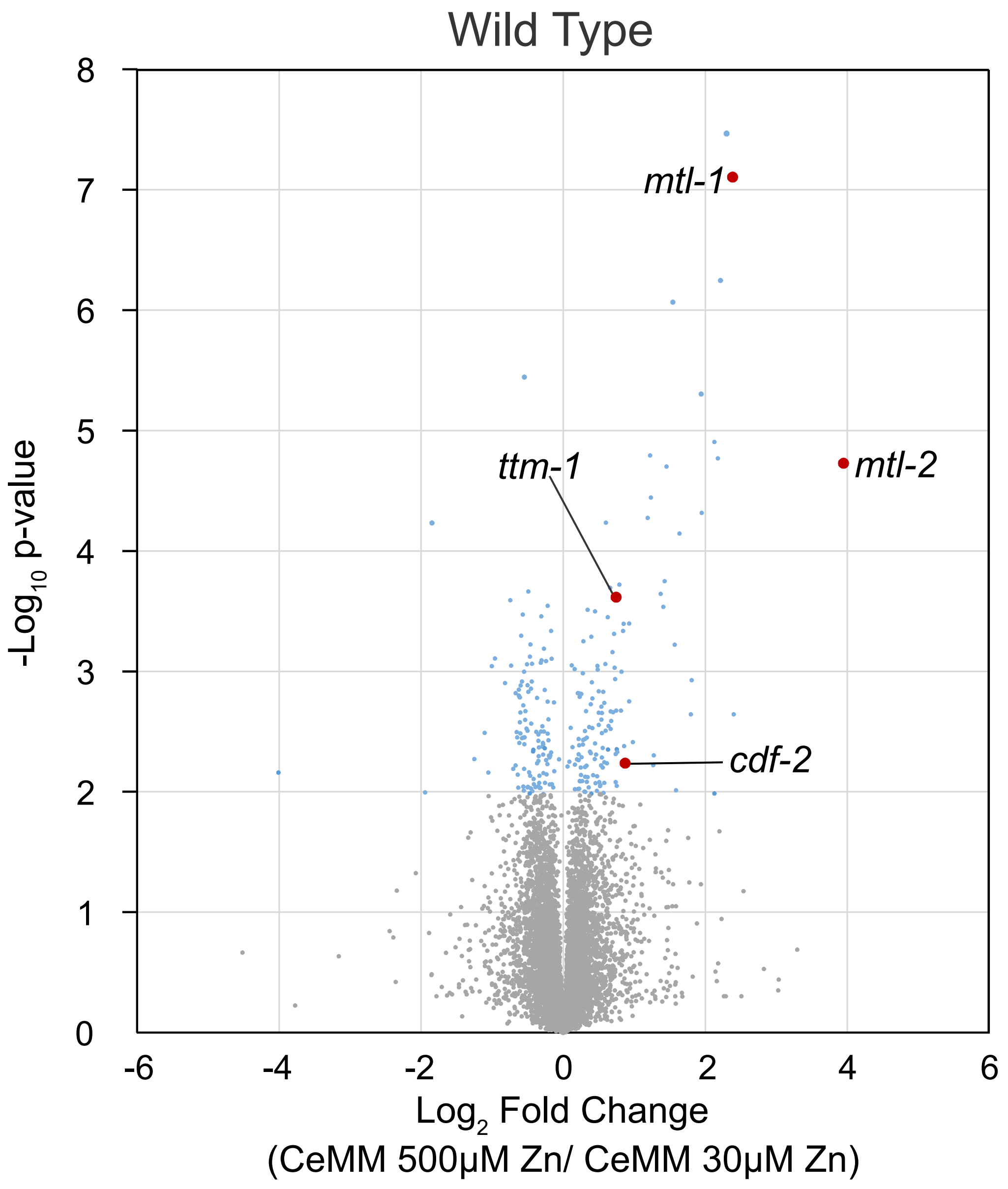
Wild Type
8
7
6
5
4
3
2
1
0
-Log10 p-value
-6
-4
-2
0
2
4
6
Log2 Fold Change
(CeMM 500μM Zn/ CeMM 30μM Zn)
mtl-1
mtl-2
ttm-1
cdf-2

### Figure S2

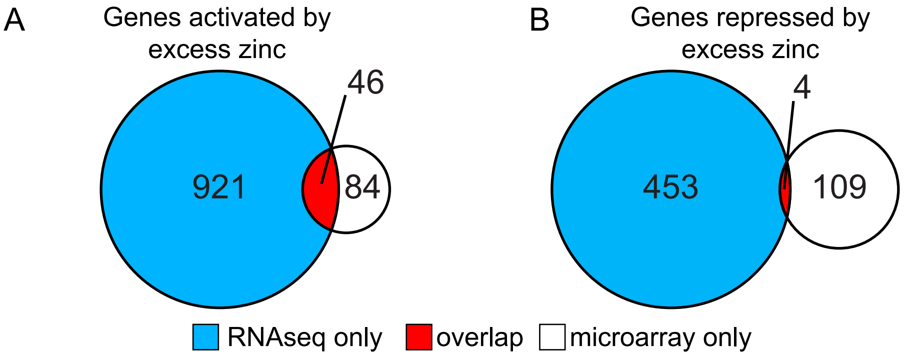


C

| | RNAseq | | Microarray | | qPCR | |
|---|---|---|---|---|---|---|
| Gene | Log$_2$ Fold Change | *p*-value | Log$_2$ Fold Change | *p*-value | Log$_2$ Fold Change | *p*-value |
| *asp-17* | 6.95 | 7.34E-14 | 2.22 | 5.67E-07 | 7.50 | <0.0001 |
| *mtl-1* | 6.92 | 3.20E-27 | 2.39 | 7.89E-08 | 7.16 | <0.0001 |
| *gst-33* | 5.83 | 1.99E-15 | 1.81 | 1.18E-03 | 5.07 | <0.0001 |
| *mtl-2* | 4.45 | 8.73E-21 | 3.95 | 1.87E-05 | 6.46 | <0.0001 |
| *cdf-2* | 3.91 | 2.34E-21 | 0.87 | 5.81E-03 | 5.80 | <0.0001 |
| *B0024.4* | 3.82 | 2.25E-07 | 1.43 | 1.79E-04 | N.D. | |
| *cpr-1* | 3.63 | 5.53E-11 | 1.37 | 2.27E-04 | 3.80 | 0.0002 |
| *thn-2* | 3.4 | 1.04E-18 | 1.22 | 1.61E-05 | 3.50 | <0.0001 |
| *spp-11* | 3.33 | 6.56E-04 | 1.57 | 6.03E-04 | N.D. | |
| *F54B8.4* | 3.08 | 1.11E-13 | 0.33 | 8.29E-03 | N.D. | |
| *ttm-1* | 3.07 | 1.49E-21 | 0.75 | 2.43E-04 | 3.73 | <0.0001 |
| *B0507.8* | 3.02 | 2.62E-07 | 0.75 | 4.44E-03 | N.D. | |
| *T01D3.6* | 2.62 | 2.32E-07 | 0.72 | 9.35E-04 | N.D. | |
| *lro-1* | 2.57 | 1.15E-08 | 0.45 | 3.17E-04 | N.D. | |
| *srlf-35* | 2.57 | 1.34E-11 | 0.56 | 1.48E-03 | N.D. | |
| *cyp-33C5* | 2.48 | 4.73E-08 | 0.17 | 9.55E-03 | N.D. | |
| *lys-7* | 2.43 | 5.41E-06 | 0.55 | 6.30E-03 | N.D. | |
| *C29F3.7* | 2.26 | 3.05E-18 | 1.54 | 8.60E-07 | N.D. | |
| *K05B2.4* | 2.23 | 2.29E-09 | 0.67 | 2.16E-03 | N.D. | |
| *C29F7.2* | 2.17 | 2.54E-17 | 0.73 | 1.16E-03 | 2.82 | <0.0001 |
| *W07B8.4* | 2.07 | 2.86E-04 | 0.93 | 4.02E-04 | N.D. | |
| *F53A9.1* | 2.04 | 7.71E-06 | 0.23 | 1.53E-03 | N.D. | |
| *clec-52* | 1.81 | 1.84E-06 | 1.45 | 1.99E-05 | 4.79 | <0.0001 |
| *cyp-33C4* | 1.77 | 1.72E-07 | 0.6 | 5.84E-05 | 2.74 | <0.0001 |
| *K08D8.6* | 1.56 | 1.76E-13 | 0.59 | 8.68E-04 | N.D. | |
| *ncx-7* | 1.49 | 2.46E-10 | 0.98 | 3.87E-03 | 2.04 | <0.0001 |
| *R05H10.1* | 1.33 | 3.41E-04 | 0.56 | 4.32E-03 | N.D. | |
| *Y46G5A.20* | 1.31 | 5.86E-04 | 0.37 | 2.91E-03 | N.D. | |
| *R09F10.1* | 1.23 | 9.31E-10 | 0.28 | 4.05E-03 | N.D. | |
| *T07D3.4* | 1.19 | 3.76E-06 | 0.7 | 2.18E-03 | N.D. | |
| *npa-1* | 1.17 | 4.23E-06 | 0.39 | 4.79E-03 | 1.77 | <0.0001 |
| *tth-1* | 1.13 | 3.82E-08 | 0.16 | 9.59E-04 | N.D. | |
| *T21H3.1* | 1.1 | 1.15E-12 | 0.6 | 3.11E-03 | N.D. | |
| *clec-83* | 1.07 | 2.46E-07 | 0.34 | 3.08E-04 | N.D. | |
| *ugt-51* | 1.01 | 2.07E-04 | 0.82 | 1.01E-03 | N.D. | |
| *clec-66* | 0.99 | 3.27E-08 | 0.79 | 1.91E-04 | N.D. | |
| *apy-1* | 0.98 | 1.32E-09 | 0.54 | 1.96E-03 | N.D. | |
| *lys-1* | 0.97 | 5.91E-10 | 0.5 | 2.21E-03 | N.D. | |
| *lys-2* | 0.96 | 1.20E-08 | 0.26 | 1.54E-03 | N.D. | |
| *Y43C5A.3* | 0.93 | 1.57E-05 | 0.39 | 1.88E-03 | N.D. | |
| *asm-2* | 0.87 | 1.98E-05 | 1.19 | 5.31E-05 | N.D. | |
| *scl-5* | 0.85 | 4.58E-04 | 0.33 | 6.09E-03 | N.D. | |
| *tmem-135* | 0.83 | 8.84E-06 | 0.81 | 2.12E-03 | N.D. | |
| *T02C5.1* | 0.69 | 4.29E-06 | 0.33 | 3.56E-03 | N.D. | |
| *ctsa-1.2* | 0.62 | 1.24E-06 | 0.36 | 9.19E-03 | N.D. | |
| *W01A11.1* | 0.47 | 9.39E-04 | 0.31 | 3.12E-03 | N.D. | |
| *acp-6* | -0.55 | 1.37E-05 | -0.31 | 3.17E-03 | N.D. | |
| *F44E7.5* | -0.63 | 5.75E-05 | -0.61 | 3.27E-03 | N.D. | |
| *C55B7.3* | -0.88 | 4.35E-04 | -1.25 | 5.34E-03 | N.D. | |
| *C14F11.4* | -0.88 | 3.61E-04 | -0.56 | 1.91E-03 | N.D. | |

### Figure S3

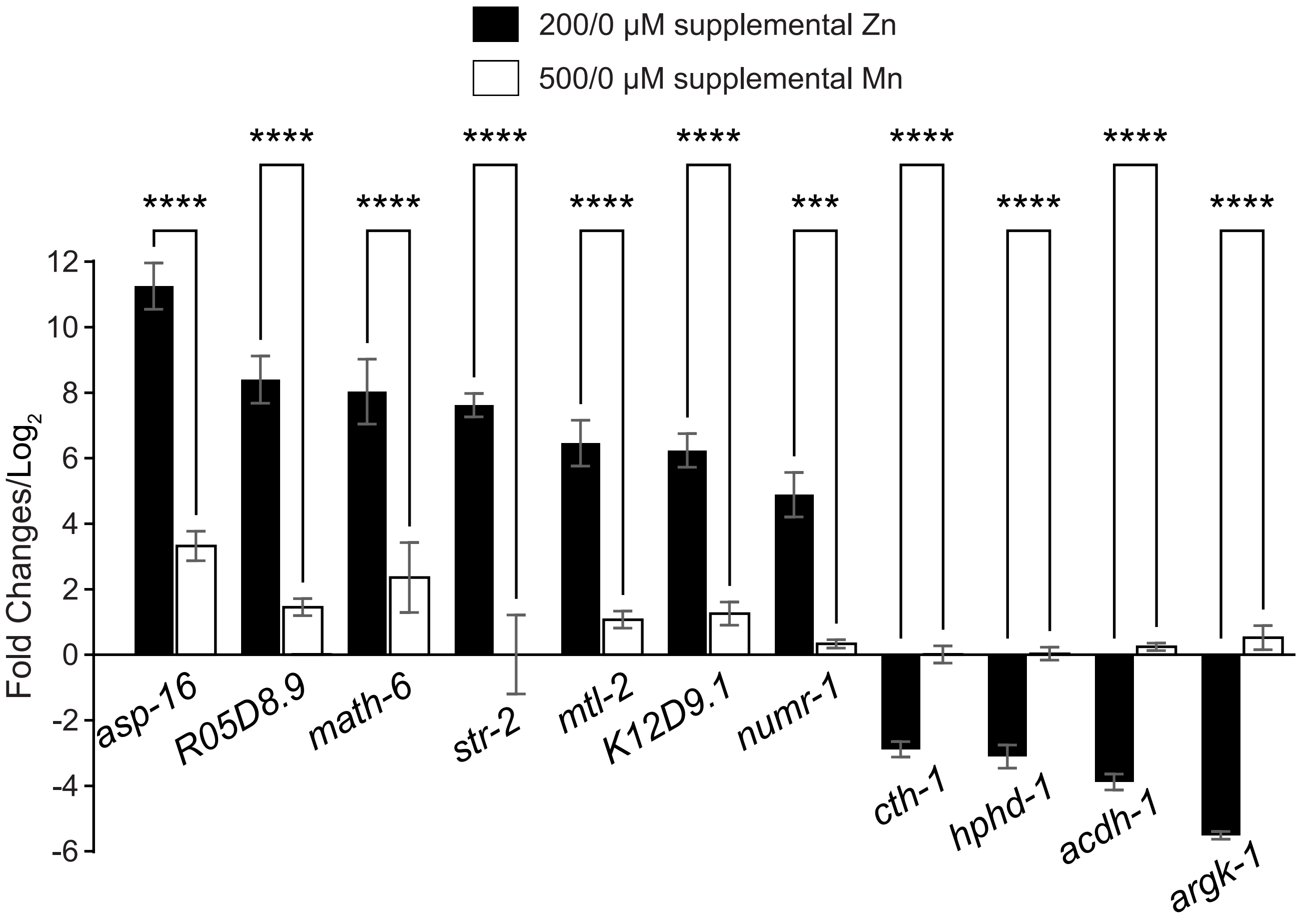
200/0 μM supplemental Zn
500/0 μM supplemental Mn
****
****
****
****
****
****
***
****
****
****
****
12
10
8
6
4
2
0
-2
-4
-6
Fold Changes/Log2
asp-16
R05D8.9
math-6
str-2
mtl-2
K12D9.1
numr-1
cth-1
hphd-1
acdh-1
argk-1

### Figure S4

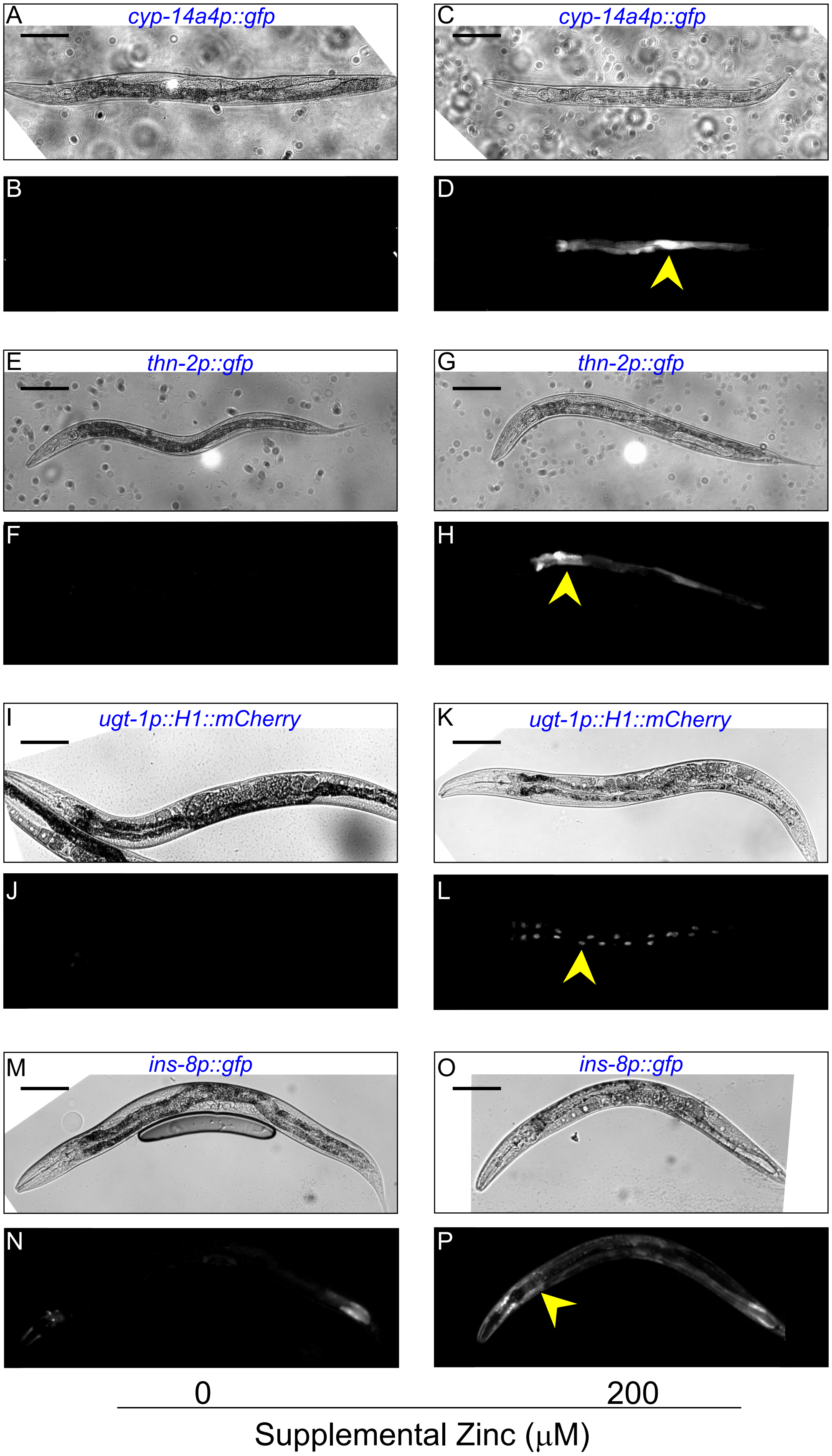

A
cyp-14a4p::gfp
C
cyp-14a4p::gfp
B
D
E
thn-2p::gfp
G
thn-2p::gfp
F
H
I
ugt-1p::H1::mCherry
K
ugt-1p::H1::mCherry
J
L
M
ins-8p::gfp
O
ins-8p::gfp
N
P
0
200
Supplemental Zinc (μM)

### Figure S5

A

| Gene | Position | E-box Sequence | *p*-value |
|---|---|---|---|
| *hlh-30* | -1165 | ACACGCGA | 6.60E-04 |
| ***hizr-1*** | **-563** | A**CACG**G**G**A | 2.40E 04 |
| *lipl-1* | -439 | CCACGTGG | 5.90E-04 |
| *lipl-2* | -55 | ACACGTGT | 9.00E-05 |
| *lipl-3* | -431 | ACACGTGC | 4.50E-04 |
| *sqst-3* | -175 | TCACGTGG | 3.10E-05 |

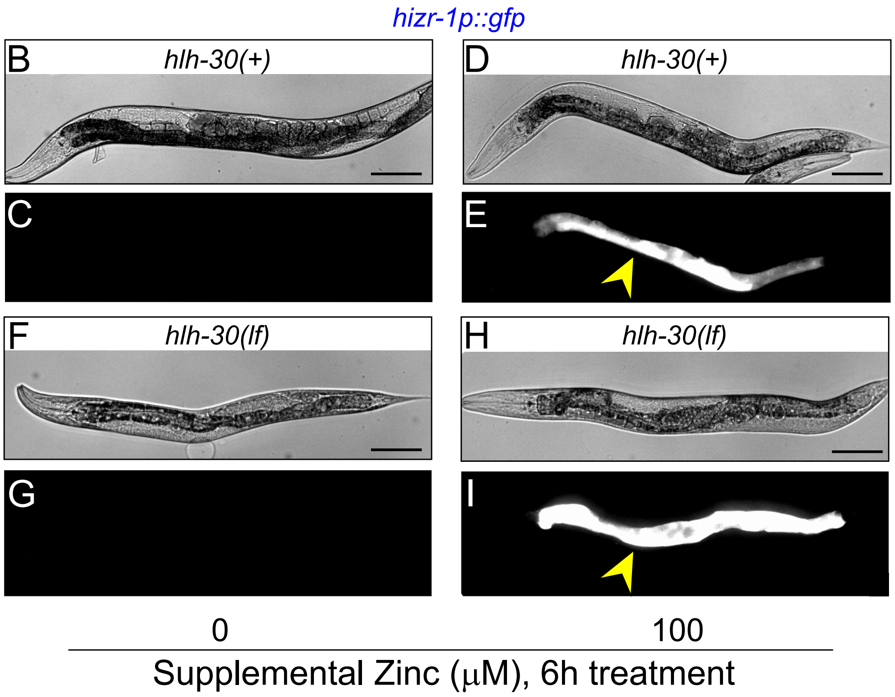
